# Mapping and longitudinal imaging of potentiated synapses in the mouse hippocampus using SynActive-eGRASP

**DOI:** 10.64898/2026.10.30.762848

**Authors:** Francesca Chiara Latini, Ajesh Jacob, Hadi Mirzapourdelavar, Antonino Cattaneo, Alessio Attardo, Marco Mainardi

**Author notes:** Equal contribution as co-first authors. Equal contribution as co-senior authors (Alessio Attardo, Marco Mainardi). Antonino Cattaneo passed away on June 28, 2026.

## Abstract

Identifying individual synapses that undergo long-term potentiation following learning at specific circuits remains a major challenge in neuroscience. To address this problem, we developed SA-eGRASP, a genetically encoded reporter that combines SynActive (SA) targeting with enhanced GFP Reconstitution Across Synaptic Partners (eGRASP). SA drives the expression of the postsynaptic half of eGRASP at potentiated spines via the 5′ and 3′ untranslated regions (UTRs) of the immediate-early gene *Arc*, thus enabling selective labeling of individual synapses that are both structurally connected and have undergone potentiation through transsynaptic GFP reconstitution. Here, we validated SA-eGRASP and combined it with high-resolution optical imaging to generate a detailed map of hippocampal CA3–CA1 synapses subjected to potentiation following contextual fear conditioning in mice. The high signal-to-noise ratio of SA-eGRASP, allowed us to track individual potentiated dendritic spines over the course of several days, to investigate the long-term dynamics of CA3–CA1 potentiated dendritic spines, and to study the selective potentiation of a subset of spines upon experience. SA-eGRASP paves the way to the investigation of synaptic engrams by enabling longitudinal, circuit-specific investigation of synapses potentiated during learning in mice performing naturalistic behaviors.

## INTRODUCTION

Elucidating the mechanisms underlying memory formation and retention has long been a central goal in neuroscience, with mounting evidence implicating synaptic connectivity and plasticity as the fundamental substrates^1–3^. While neuronal ensemble–level plasticity has been extensively studied by taking advantage of Immediate-Early Genes (IEGs) to selectively tag neurons activated during plasticity-related events^4–10^, our understanding at the synaptic level is still incomplete. While significant progress has been made to identify and track synapses between IEG-expressing neurons^11–14^, identifying the individual synapses that undergo potentiation during learning and memory processes remains a major challenge^5,15^. We previously tackled this problem by devising SynActive (SA), a translation-dependent strategy that exploits the 5′ and 3′ untranslated regions (UTRs) of *Arc* mRNA, which confer dendritic targeting and synapse-specific translational control to drive activity-dependent reporter expression at potentiated dendritic spines. Coupled with postsynaptic targeting peptide motifs, SA selectively labels recently potentiated synapses with high spatial specificity^16,17^. However, two key open questions remain: (i) how potentiated synapses between defined neuronal populations during a defined learning and memory process can be selectively identified, and (ii) how these potentiated synapses evolve over time *in vivo* during memory formation and retention.

To address these questions, we generated SA-eGRASP, a synaptic potentiation–dependent eGRASP^13^ reporter (SA-eGRASP). As classic eGRASP, SA-eGRASP is based on the reassembly of split GFP fragments expressed on the two sides of the synaptic cleft^13,18,19^. However, in SA-eGRASP, the postsynaptic split GFP moiety is expressed under SynActive control^16,17,20^, thereby restricting the reconstitution of a functional GFP at potentiated synapses. We first characterized SA-eGRASP expression in neuronal cultures, then applied it to the hippocampal CA3–CA1 pathway to identify potentiated connections between defined pre- and postsynaptic neuronal populations. We exploited SA-eGRASP to generate a high-resolution cartography of synapses undergoing potentiation throughout various stages of contextual fear conditioning (CFC). Finally, we tracked the longitudinal dynamics of identified synapses potentiated upon exposure to an enriched environment (EE) *in vivo*, thus characterizing the long-term temporal dynamics of active synapses in behaving mice during naturalistic conditions.

## MATERIALS AND METHODS

### Gene constructs

Constitutive post-eGRASP (Addgene #111584) was previously described in Choi et al.^13^. SA-post-eGRASP was generated by sequentially cloning the TRE3g promoter, Arc 5′ untranslated region (UTR), myr-mScarlet-P2A-post-eGRASP, Arc 3′ UTR, and an SV40 short polyadenylation signal (spA) into pAAV-hSyn::EGFP-WPRE-pA (Addgene #50465) between the inverted terminal repeats (ITRs). TRE3g, Arc 5′ UTR and Arc 3′ UTR sequences corresponded to nucleotides 1–230 and 1424–3026 of the rat Arc transcript (NM_019361.2) and were derived from Gobbo et al.^16^. Unlike the original SynActive construct, SynTag was omitted because synaptic targeting was provided by the neuroligin stalk that is integral part of the post-eGRASP moiety. The spA sequence was derived from pAAV-CW3SL-EGFP (Addgene #61463).

For pre-eGRASP expression, pAAV-hSyn::EGFP-WPRE-pA (Addgene #50465) was replaced with TRE3g and pre-eGRASP (Addgene #111598) to generate pAAV-TRE3g::pre-eGRASP-WPRE-pA.

Driver constructs were generated by cloning rtTA-P2A and tdTomato or mTurquoise2 between the hSyn promoter and WPRE to obtain hSyn::rtTA-P2A-tdTomato and hSyn::rtTA-P2A-mTurquoise2. rtTA2S-M2 was amplified from Gobbo et al.^16^, and a P2A sequence was added by overhang PCR. All plasmids were constructed by VectorBuilder and Addgene. For *in vivo* transduction, pAAV-TRE3g::pre-eGRASP, pAAV-TRE3g::post-eGRASP, pAAV-SA-post-eGRASP, pAAV-hsyn::rtTA-P2A-tdTomato, and pAAV-hsyn::-rtTA-P2A-mTurquoise2 were packaged into adeno-associated virus (AAV) particles belonging to the PHP.eB serotype using standard in vitro procedures performed by Innovavector (Naples, Italy), yielding preparations with a titer ranging from 1.9×10^12^ to 6.1×10^12^ vg/ml.

### Primary neuronal culture, transfection, and treatments

All procedures involving primary neuronal cultures were approved by the Italian Ministry of Health and complied with European regulations governing animal research (authorization 65E5B.N.ZDO).

Primary hippocampal cultures were prepared from postnatal day (P) 0 B6/129 mice as described previously^17^. Neurons were transfected using Lipofectamine 2000 (Invitrogen) according to the manufacturer’s instructions.

For *in vitro* SynActive-eGRASP experiments, two independent transfection mixtures were prepared. The presynaptic mixture contained Const-pre-eGRASP (0.5 μg) and tdTomato filler (0.5 μg) diluted in 25 μl Opti-MEM and mixed with 1.5 μl Lipofectamine 2000 diluted in 25 μl Opti-MEM. The postsynaptic mixture contained either SynActive-post-eGRASP or Const-post-eGRASP (0.5 μg) together with mTurquoise2 filler (0.5 μg), prepared identically. Following a 5-min incubation, both mixtures were added to the same well and incubated for 2.5 h at 37°C. The transfection medium was then replaced with 350 μl conditioned medium and 150 μl fresh neuronal growth medium. For single-component controls, filler DNA was increased to 1 μg. For AMPAR and NMDAR enrichment experiments, tdTomato and mTurquoise2 fillers were used alongside the corresponding pre- and postsynaptic eGRASP components.

Chemical LTP was induced as described by Lu et al.^53^. Conditioned medium was collected before treatment. Neurons were incubated for 30 min at 37°C in extracellular solution containing 140 mM NaCl, 1.3 mM CaCl2, 5 mM KCl, 25 mM HEPES, 33 mM D-glucose, 0.5 μM tetrodotoxin (TTX, Tocris BioSciences 1078), 1 μM strychnine (Sigma-Aldrich S0532), and 50 μM picrotoxin (PTX, Tocris BioSciences 1128). The solution was replaced with an extracellular solution containing 200 μM glycine for 3 min, then returned to the normal extracellular solution for 30 min. The conditioned medium was then restored, and neurons were maintained until fixation 24 h later. For NMDAR blockade experiments, 50 μM D-AP5 (Sigma-Aldrich A8054) was applied together with doxycycline, and cultures were fixed after 36 h.

### *In vitro* immunofluorescence

Neurons were fixed as previously described^17^. Cultures were blocked in 5% BSA in PBS for 1 h and incubated with primary antibodies in 2.5% BSA in PBS for 2-3 h. Primary antibodies included PSD-95 (Synaptic Systems 124011, 1:500), Synaptophysin-1 (Synaptic Systems 101011, 1:500), GluA1 (Cell Signaling 13185, 1:500), and GluN2A/B (Synaptic Systems 244003, 1:500). Following washing in PBS, cultures were incubated for 1 h with Alexa Fluor 647-conjugated anti-rabbit (Thermofisher A-31573) or anti-mouse secondary antibodies (Thermofisher A-31571) diluted in 2.5% BSA in PBS. Samples were washed and mounted for imaging.

### *In vitro* imaging and analysis

Confocal z-stacks (1024×1024 pixels; 0.4 μm step size) were acquired using either Leica SP2, Leica SP5, or Zeiss LSM800 confocal microscopes equipped with high numerical aperture oil immersion objectives (HCX PL APO 63×/1.4 numerical aperture (NA), and Plan-Apochromat 63×/1.4 NA DIC M27, respectively). Sequential excitation was used to minimize spectral overlap.

A synapse was defined as an overlap or close apposition between a tdTomato+ presynaptic bouton and an mTurquoise2+ dendrite. Synapses were counted manually using the ImageJ Cell Counter plugin. Neurons with fewer than 10 identified synapses were excluded.

For colocalization analyses, line ROIs extending from spine heads to dendritic shafts were drawn and intensity profiles extracted from maximum projections. Profiles were normalized by subtracting the minimum intensity and dividing by the intensity range. Average profiles were generated after alignment to the maximum green fluorescence signal. Colocalization percentages were calculated as the proportion of eGRASP puncta overlapping PSD-95 or apposed to Synaptophysin-1.

For receptor enrichment analyses, circular ROIs corresponding to spine heads were drawn over eGRASP-positive spines. Background-subtracted integrated GluA1 or GluN2A/B fluorescence was measured. Spine volume was estimated as 4/3*πr*^3^, where *r* is the radius of the corresponding spine ROI.

### Animals

C57BL/6J mice aged 3–4 or 2-3 months were used for *ex vivo* and *in vivo* experiments, respectively. Animals were housed in groups of up to four under a 12 h light/dark cycle with ad libitum access to food and water. Experiments were conducted during the light phase. *Ex vivo* procedures were performed under authorization B4BB8.20.EXT.10 by the Italian Ministry of Health. *In vivo* procedures complied with Leibniz Association guidelines and were approved under license 42502-2-1737 LIN.

### Stereotaxic injections

Mice were anesthetized using zoletil/xylazine (80/10 mg/kg). Body temperature was maintained at 36°C using a closed-loop heating system. Craniotomies were made above right CA3 (AP −1.75 mm, ML 2.35 mm to bregma; DV −1.675 mm) and left CA1 (AP −1.80 mm, ML −1.5 mm to bregma; DV −1.3 mm)^54^. Viral suspensions (0.25–0.5 μL) were delivered at 0.1 μl/min through heat-pulled, calibrated glass micropipettes (Sigma-Aldrich Z114952) using an oil injector (CellTram Oil, Eppendorf, Germany).

For *ex vivo* experiments, the CA3 injection site received a mix of hSyn-rtTA-P2A-mTurquoise2 (3.05×10^12^ vg/ml) and pre-eGRASP AAVs (0.95×10^12^ vg/ml). The contralateral CA1 injection site received a mix of hSyn-rtTA-P2A-tdTomato filler (0.74×10^12^ vg/ml) and SA-post-eGRASP (1.52×10^12^ vg/ml) AAVs.

Pipettes remained in place for 10 min following injection before withdrawal. Incisions were sutured with Vicryl 5/0 non-absorbable suture, and mice received postoperative analgesia using oral paracetamol (100 mg/kg) or Metacam (0.5 mg/ml) for 3–5 d.

For *in vivo*, two-photon imaging experiments, a combination of AAV1/2-hSyn::rtTA (3.2×10^13^ vg/ml), AAV-PHP.eB-TRE3g::preSA-eGRASP (1.9×10^12^ vg/ml) was injected into the dCA3, and AAV1/2-CamKII::Cre (2.3×10^12^ vg/ml), AAV1/2-DIO-tdTomato-P2A-rtTA (3.6×10^14^ vg/ml) and AAV-PHP.eB-TRE3g::myrScarlet-P2A-postSA-eGRASP (1.9×10^12^ vg/ml) in the contralateral dCA1.

### Contextual Fear Conditioning (CFC) and doxycycline labeling

Behavioral experiments were conducted 21–28 d after AAV injections using a sound-insulated testing chamber (#46001, Ugo Basile, Italy). Mice were habituated to the conditioning chamber for 5 min. Twenty-four hours later, mice underwent the encoding phase of the conditioning memory paradigm, consisting of five 0.5-mA foot shocks (1 s duration) delivered at 25 s intervals after an initial 30-s exploration period. Animals remained in the chamber for an additional 2 min after the final shock. Memory recall was assessed 24 h later by re-exposure to the same context for 5 min. Freezing was quantified automatically using AnyMaze.

Gene construct expression windows were opened using two intraperitoneal injections of doxycycline (150 μL, 10 mg/ml) separated by 20 h. The doxycycline injection schedules for each experimental group are indicated in Fig. 3a.

For activity induction independent of fear conditioning, mice received kainic acid (5 mg/kg, i.p.) in approximately 0.2 ml saline. Doxycycline administration and subsequent procedures remained unchanged.

### SA-eGRASP signal following local D-AP5 infusion

For D-AP5 infusion experiments, mice received unilateral stereotaxic viral injections as described above. Sixteen days later, custom 26G guide cannulae (D/V −1.3 mm) were implanted above CA1 in the hemisphere expressing SA-post-eGRASP and secured with dental cement. Following at least 5 days of recovery, mice were anesthetized with isoflurane and received local CA1 infusions of either pH-adjusted D-AP5 (40 mM) or filtered saline (final volume 1 μl at 0.1 μl/min). The injection needle was left in place for 1 additional minute before withdrawal. Infusions were performed 15 min before the association phase of contextual fear conditioning. Mice were transcardially perfused 3 h after completion of the CFC association phase.

### Tissue processing and confocal imaging

Animals received terminal anesthesia with an overdose of chloral hydrate (10% w/v in saline) and perfused transcardially with PBS followed by 4% PFA in 0.1 M phosphate buffer (PB), pH 7.4. Brains were post-fixed overnight and cryoprotected in 30% sucrose in PB. Unless otherwise specified, animals were perfused 24 h after the labeling session.

Coronal sections (80 μm) were collected using a sliding microtome and mounted in VectaShield (Vector Labs, USA). The CA1 area was imaged using a Leica SP2 confocal microscope equipped with a 63×/1.4 NA objective. Images were acquired at a voxel size of 0.116×0.116×0.396 μm. Four overlapping image tiles spanning all CA1 layers were acquired per section.

### Analysis of confocal microscopy images

*SA-eGRASP signal intensity analysis.* The raw integrated density of SA-eGRASP expression levels was measured from maximum intensity projections acquired using a Zeiss LSM900 microscope, using ImageJ. Thresholds were determined from background ROIs, and raw integrated density was calculated above threshold.

*Image stitching.* Image tiles were stitched using the BigStitcher plugin in ImageJ. Following manual alignment, intensity-based optimization and image fusion were performed. Background subtraction was applied using a rolling-ball algorithm before RGB stack generation.

*Tracing of dendritic segments.* Dendritic segments in stratum radiatum were reconstructed in NeuTube and exported as SWC files. Traces were resampled and standardized in Vaa3D. *Automated SynActive-eGRASP analysis.* Analyses were conducted using customized MATLAB scripts. As tdTomato expression in CA1 dendrites was unsuitable for reliable spine volume quantification, image analysis was limited to the channels expressing SA-eGRASP and mTurquiose2. First, SA-eGRASP and mTurquoise2 channels from the stitched RGB image were loaded into the program. Although SA-eGRASP puncta were also observed in the SO, analyses were not performed due to limited fidelity in tracing thin, densely labelled basal dendrites. Pixel intensity values were then corrected for photobleaching along the z-axis by multiplying by a bleach ratio 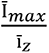 ^55^. The corrected pixel value is given by 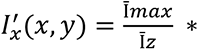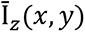, where Īz represents the average blue intensity calculated for each z-slice, and Ī_max_ denotes the maximum Ī_z_ value. To delimit the analysis, a mask was then created around each trace, corresponding to a 31×31×9-pixel cuboid (equivalent to 3.596×3.596×3.564 μm in x, y, and z dimensions). Detection algorithms described in Feng, Zhao and Kim^56^, were used to automatically identify SA-eGRASP signals, corresponding to potentiated CA3-CA1 synapses, and mTurquoise2 signals from CA3 presynaptic boutons. Parameters for puncta detection (I_min_, V_max_, and V_min_) were empirically determined to minimize false positives and negatives. V_min_ values used for SA-eGRASP and mTurquoise2 were 25 pixels and 50 pixels, respectively, while V_max_ was set at 115 pixels for both. While a constant I_min_ value of 30 was applied for SA-eGRASP, I_min_ values for mTurquoise2 showed slight variations. Manual corrections were then applied for over-segmentation errors, involving only deletions in the case of SA-eGRASP, and joining or deleting detected mTurquoise2 puncta.

As a further level of stringency, an SA-eGRASP punctum was considered a true positive only if the Euclidean distance between its centroid and the centroid of its nearest neighbor mTurquoise2 punctum was less than 2 μm. SA-eGRASP density and percentage were calculated by dividing the number of SA-eGRASP puncta by the pathlength of the dendritic trace (measured using neuTube) and the number of mTurquoise2 puncta, respectively. CA3-CA1 synaptic density is given by the ratio of detected mTurquoise2 puncta to the dendritic trace pathlength. The volume of SA-eGRASP puncta is calculated by multiplying the number of voxels in a detected punctum by the voxel volume.

The distribution of detected puncta along the stratum radiatum was determined by first mapping the mTurquoise2 puncta to the closest node in the corresponding dendritic trace, followed by mapping the SA-eGRASP puncta to the trace node of their colocalized mTurquoise2 puncta. This resulted in two groups of trace nodes: those with mTurquoise2 puncta only and those with colocalized mTurquoise2 and SA-eGRASP puncta. The relative position (*di*) of the trace node in the stratum radiatum was calculated as 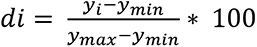 where y_i_, y_min_ and y_max_ represent the y-coordinate of the mapped node i.e., the start and the end of the *stratum radiatum*, respectively. Nodes were then binned according to their *di* values, where the bin-width of 10% approximately corresponds to 20 μm. The percentage of SA-eGRASP nodes, the density of mTurquoise2, and the volume of SA-eGRASP puncta in each bin were calculated.

To perform clustering analyses, pathlength distances were calculated for each dendritic node, with colocalized SA-eGRASP puncta, to the branching node (or root node if no branches were present), corresponding to a reduction in the spatial dimension from 3D (x, y, z) to 1D (pathlength). Frequentative summation of the Euclidean distance between two adjacent nodes, starting from the SA-eGRASP node and ending in the branching node, was used to calculate pathlength. The distance from one SynActive-eGRASP node to another within the branch was determined by subtracting their respective path lengths from the branching node. The minimum value of these distances gave its nearest neighbor distance (d). Clustering analysis was performed using the MATLAB function - *dbscan* with parameters, input matrix – SA-eGRASP node pathlength from the branch, epsilon neighborhood – median dCFC, and minimum number of points – 3^57^. This defined a “cluster” as a group of three or more SA-eGRASP-positive puncta with nearest neighbor distances smaller than the median nearest neighbor distance observed in the HC group (median dHC, 1.16 μm). Cluster density was calculated by normalizing the number of clusters in a trace to its total path length.

The customized codes utilized in this analysis, along with a detailed user manual, are available through GitHub upon request.

### Optical cannula implantation and longitudinal two-photon imaging

Optical cannulae were implanted as described previously^58^. After craniotomy and cortical aspiration, a coverslip-cannula assembly was positioned above the hippocampus and secured using dental acrylic and a custom head plate. Mice recovered for at least two weeks before imaging.

Expression was induced by doxycycline injection, and animals were imaged for 14 days under low-dose isoflurane anesthesia. Imaging was performed on a Thorlabs Bergamo microscope equipped with 920 nm and 1040 nm lasers and a 40×/0.8 NA water-immersion objective. Image stacks were acquired at 0.1079×0.1079×0.5 μm voxel size with 50–100 z-planes and ten repeated acquisitions per plane.

### Two-Photon Image Processing and Spine Tracking

Repeated acquisitions were rigidly registered to average images using custom Python scripts and subsequently averaged. Stacks were deconvolved using AutoQuantX3, depth-normalized, and temporally registered using cross-correlation.

Spines were tracked manually using TrackMate^59^. Each spine was assigned a unique identifier based on its coordinates. Reappearing spines were assigned new identifiers. Spines exhibiting only red fluorescence were classified as SA-eGRASP^-^, whereas those containing green fluorescence were classified as SA-eGRASP+.

### Statistical Analysis

Statistical analyses were performed using GraphPad Prism (version 9; GraphPad Software) unless otherwise stated. Linear mixed-effects models were fitted in R (version 4.6.0) using the nlme package (version 3.1-169). All statistical tests were two-sided. *P* < 0.05 was considered statistically significant.

Unless otherwise indicated, data are presented as mean ± s.e.m., and each data point represents the mean value obtained from an individual mouse. Statistical analyses were performed on mouse-level means unless otherwise specified in the figure legends. For analyses in which individual dendritic traces, neurons, spines, puncta or clusters constituted the statistical unit, the corresponding sample sizes and units of analysis are indicated in the relevant figure legends. Exact sample sizes (N), representing independent biological replicates, are reported for each experiment.

Comparisons between two groups were performed using paired or unpaired, two-tailed Student’s t-tests, as appropriate. Comparisons among three or more groups were performed using one-way ANOVA (ANOVA-1), two-way ANOVA (ANOVA-2) or repeated-measures ANOVA, followed by Tukey’s, Holm–Šídák’s or Fisher’s least significant difference (LSD) multiple-comparisons tests, as specified in the corresponding figure legends. Relationships between behavioral freezing and SA-eGRASP measurements were assessed using simple linear regression. Cumulative distributions were compared using two-sided Kolmogorov– Smirnov tests. Survival curves were compared using the log-rank (Mantel–Cox) test.

Variability differences were assessed using linear mixed-effects models with mouse as a random effect and group-specific residual variances implemented in R (package: *nlme*). Statistical analyses accounted for the nesting of branch measurements within individual mice. Model fits were compared using likelihood-ratio tests. Equality of variances was assessed using Brown–Forsythe and Bartlett’s tests where indicated.

Exact *P* values, test statistics (including *t*, *F*, *χ²* or *D* statistics, as appropriate), degrees of freedom, regression statistics, and post-hoc multiple-comparison results are reported in the corresponding figure legends. Graphs displaying cumulative distributions report the number of observations analysed, whereas graphs summarizing biological replicates report the number of independent mice, unless otherwise stated.

### Data Availability

The original datasets generated and analyzed during the current study are available from the corresponding authors upon request.

### Code Availability

MATLAB analysis codes and guidelines are available on GitHub at https://github.com/SynActiveLab/GRASP-Analysis, and will be made public upon publication.

## RESULTS

### Generation of the SA-eGRASP constructs

To label synapses with high specificity and spatial resolution, we leveraged the reassembly of split GFP fragments expressed on the two sides of the synaptic cleft using eGRASP^13^. To ensure expression of eGRASP in synapses undergoing translation-dependent plasticity, we placed the mRNA coding for the postsynaptic eGRASP moiety under the translational control of the SA system^16^. To this aim, we generated a construct (SA-post-eGRASP) in which we inserted the postsynaptic eGRASP fragment (GFP strand 11) between the 5′ and 3′ UTR regions of the *Arc* mRNA. We placed the SA-post-eGRASP coding sequence under the transcriptional control of the tetracycline-responsive element (TRE) to achieve temporal control over its expression. Similarly, we generated a TRE-controlled version of the presynaptic moiety of eGRASP (pre-eGRASP; GFP strands 1-10). Tet-ON-dependent expression of both SA-post-eGRASP and pre-eGRASP was driven by the reverse tetracycline-dependent transactivator (rtTA, tet-ON system) under the Synapsin-1 promoter. Additionally, the constructs encoded for either mTurquoise2 or tdTomato, enabling visualization of postsynaptic dendrites and dendritic spines or presynaptic axons (Fig. 1a).

**Figure 1.**
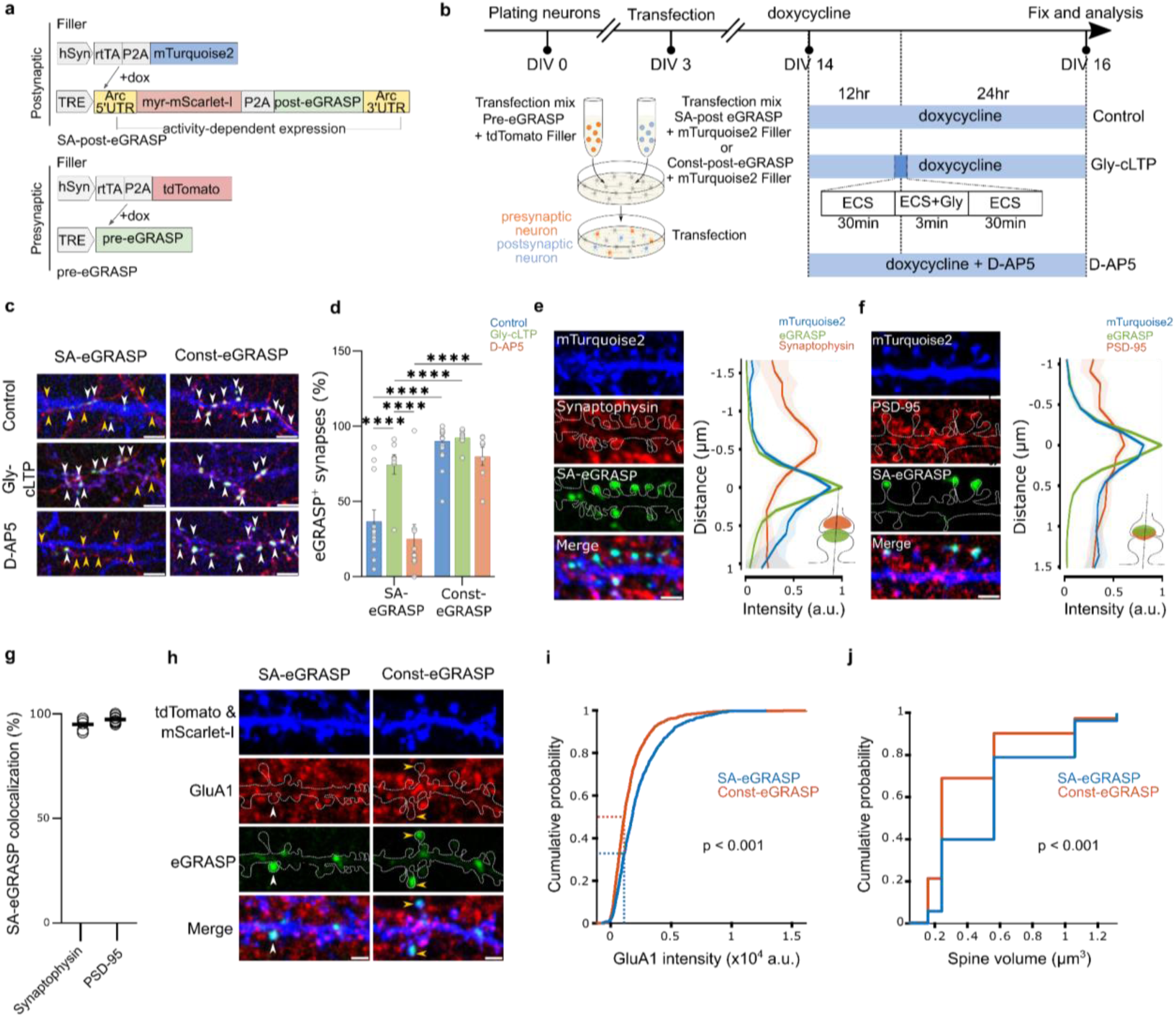
SA-eGRASP is expressed at potentiated synapses *in vitro*. **a.** Schematic description of the gene constructs used. **b.** Experimental timeline for chemical modulation of neuronal activity in co-cultured neurons expressing pre-eGRASP (red) and SA-post-eGRASP (blue). **c.** Representative confocal images of SA-eGRASP and Const-eGRASP under doxycycline (Dox)-only (Control), Dox and Glycine-mediated chemical LTP (Gly-cLTP) or Dox, Gly-cLTP and D-2-amino-5-phosphonovaleric acid (D-AP5) conditions. mTurquoise2, blue; tdTomato, red; SA-eGRASP, green. White arrowheads indicate overlapping mTurquoise2+ dendrites and tdTomato+ axons with GFP+ puncta; yellow arrowheads indicate non-overlapping contacts. Scale bar, 5 μm. **d**. Quantification of GFP-positive synapses in SA-eGRASP and Const-eGRASP neuronal cultures (ANOVA-2, treatment, group × treatment, *F*_2,122_ = 13.82, *P* < 0.0001; Tukey’s post hoc test, *P*_Control SA-eGRASP vs. Control Const-eGRASP_ < 0.0001, *P*_Control SA-eGRASP vs. Gly-cLTP SA-eGRASP_ < 0.0001, *P*_Control SA-eGRASP vs. Gly-cLTP Const-eGRASP_ < 0.0001, *P*_Control SA-eGRASP vs. D-AP5 Const-eGRASP_ < 0.0001, *P*_Control Const-eGRASP vs. Gly-cLTP SA-eGRASP_ < 0.0001, *P*_Control Const-eGRASP vs. D-AP5 SA-eGRASP_ < 0.0001, *P*_Gly-cLTP SA-eGRASP vs. Gly-cLTP Const-eGRASP_ < 0.0001, *P*_Gly-cLTP SA-eGRASP vs. D-AP5 SA-eGRASP_ < 0.0001, *P*_Gly-cLTP SA-eGRASP vs. D-AP5 Const-eGRASP_ = 0.0004, *P*_Gly-cLTP Const-eGRASP vs. D-AP5 SA-eGRASP_ < 0.0001, and *P*_D-AP5 SA-eGRASP vs. D-AP5 Const-eGRASP_ < 0.0001; SA-eGRASP: N_control_ = 23, N_Gly-cLTP_ = 22, N_D-AP5_ = 21 neurons; 4 experimental replicates; Const-eGRASP: N_control_ = 22, N_Gly-cLTP_ = 20, N_D-AP5_ = 20 neurons; 5 experimental replicates). **e, f.** Representative images and intensity profiles showing colocalization of SA-eGRASP+ puncta (green) on mTurquoise2+ dendrites with synaptophysin (**e**) or PSD-95 (**f**). Scale bar, 2 μm**. g.** SA-eGRASP colocalization with synaptophysin (N_spines_ = 856, N_neurons_ = 6) and PSD-95 (N_spines_ = 1177, N_neurons_ = 8; 2 experimental replicates). **h.** Representative images of GluA1 expression at SA-eGRASP+ (white arrowheads) and Const-eGRASP+ (yellow arrowheads) synapses. Scale bar, 2 μm. **i, j.** Cumulative distributions of GluA1 intensity (Kolmogorov–Smirnov test, *D* = 0.2074, *P* < 0.0001) and GFP+ spine volume (Kolmogorov-Smirnov Test, *D* = 0.2912, *P* < 0.0001) for SA-eGRASP and Const-eGRASP synapses (SA-eGRASP N_spines_ = 1163, Const-eGRASP N_spines_ = 1220; 2 experimental replicates).

**Extended Data Figure 1.**
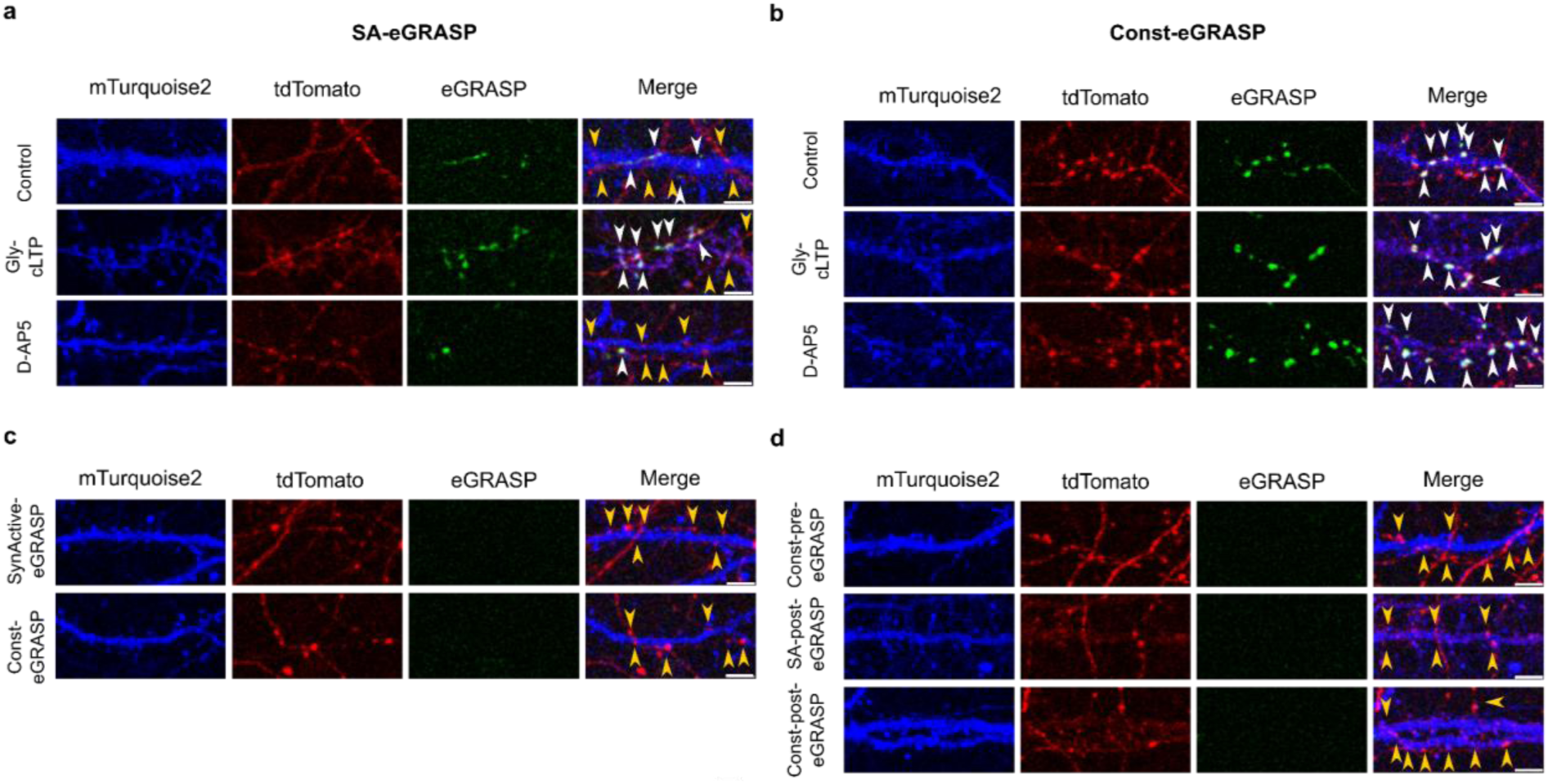
SA-eGRASP *in vitro* is not expressed in the absence of doxycycline or of one of the eGRASP components. **a,b.** Representative confocal images of SA-eGRASP (**a**) and Const-eGRASP (**b**) under doxycycline (Dox)-only (Control) conditions, Dox + Glycine-mediated chemical LTP (Gly-cLTP), and Dox + Gly-cLTP + D-2-amino-5-phosphonovaleric acid (D-AP5). White arrowheads indicate overlapping mTurquoise2+ dendrites and tdTomato+ axons with GFP+ puncta; yellow arrowheads indicate non-overlapping contacts. Scale bar, 5 μm. **c,d.** Representative images showing intersections between mTurquoise2+ dendrites and tdTomato+ axons lacking SA-eGRASP puncta (yellow arrowheads), in the absence of doxycycline (**c**) or when only one eGRASP component was expressed (**d**). Scale bar, 5 μm.

### SA-eGRASP identifies potentiated excitatory synapses in primary hippocampal neuron cultures

To ascertain that SA-eGRASP expression was synaptic potentiation-dependent, we established two sets of neuronal co-cultures. The first set contained neurons expressing either pre-eGRASP or SA–post-eGRASP, while those of the second set expressed either pre-eGRASP or the standard, SA-independent version of eGRASP (Const–post-eGRASP) (Fig. 1b). We first triggered the expression of the eGRASP constructs by adding doxycycline to the cultures, then induced chemical LTP (cLTP) or added the NMDA receptor (NMDAR) antagonist, D-AP5. Finally, we quantified the fraction of eGRASP+ synapses by confocal microscopy (Fig. 1b). Importantly, eGRASP native fluorescence could be detected directly with no need of signal amplification *via* immunofluorescence (Fig. 1c. Extended Data Fig. 1a,b). Induction of cLTP significantly increased the fraction of SA-eGRASP+ puncta at contacts between mTurquoise+ dendrites and tdTomato+ axons compared to control cultures treated with doxycycline only. Conversely, treatment with D-AP5 reduced the fraction of SA-eGRASP+ puncta with respect to controls. In contrast, in Const-eGRASP co-cultures, ∼90% of synapses displayed eGRASP+ puncta irrespective of cLTP induction or D-AP5 treatment (Fig. 1c,d). We did not detect SA-eGRASP+ puncta in the absence of doxycycline (Extended Data Fig. 1a,b) or when either pre-eGRASP or SA–post-eGRASP were omitted from the transfection mix (Extended Data Fig. 1c,d). SA-eGRASP+ spines co-localized with synaptophysin and PSD-95 (Fig. 1e-g) and exhibited significantly higher levels of the AMPA receptor (AMPAR) subunit GluA1 as well as larger spine head volumes when compared to Const-eGRASP+ spines (Fig. 1h-j).

These findings demonstrate that SA-eGRASP reports NMDAR-dependent synaptic potentiation.

### SA-eGRASP enables circuit-specific labeling of potentiated synapses

We next characterized SA-eGRASP expression *in vivo* to investigate its spatial distribution and temporal dynamics following a behavioral learning paradigm. To this aim, we targeted the hippocampal CA3–CA1 pathway, a well-characterized circuit that undergoes synaptic potentiation following CFC^21^. To express the pre- and post-synaptic constructs in distinct neuronal populations, we stereotaxically injected the dorsal (d) CA3 with adeno-associated viruses (AAVs) containing TRE-pre-eGRASP and rtTA-mTurquoise2, and the contralateral dCA1 with AAVs containing TRE-SA-post-eGRASP and rtTA-tdTomato (Fig. 2a). This ensured the SA-eGRASP fluorescent signal was genuinely intercellular and did not derive from co-expression of the pre- and post-synaptic moieties in the same neuron.

**Figure 2.**
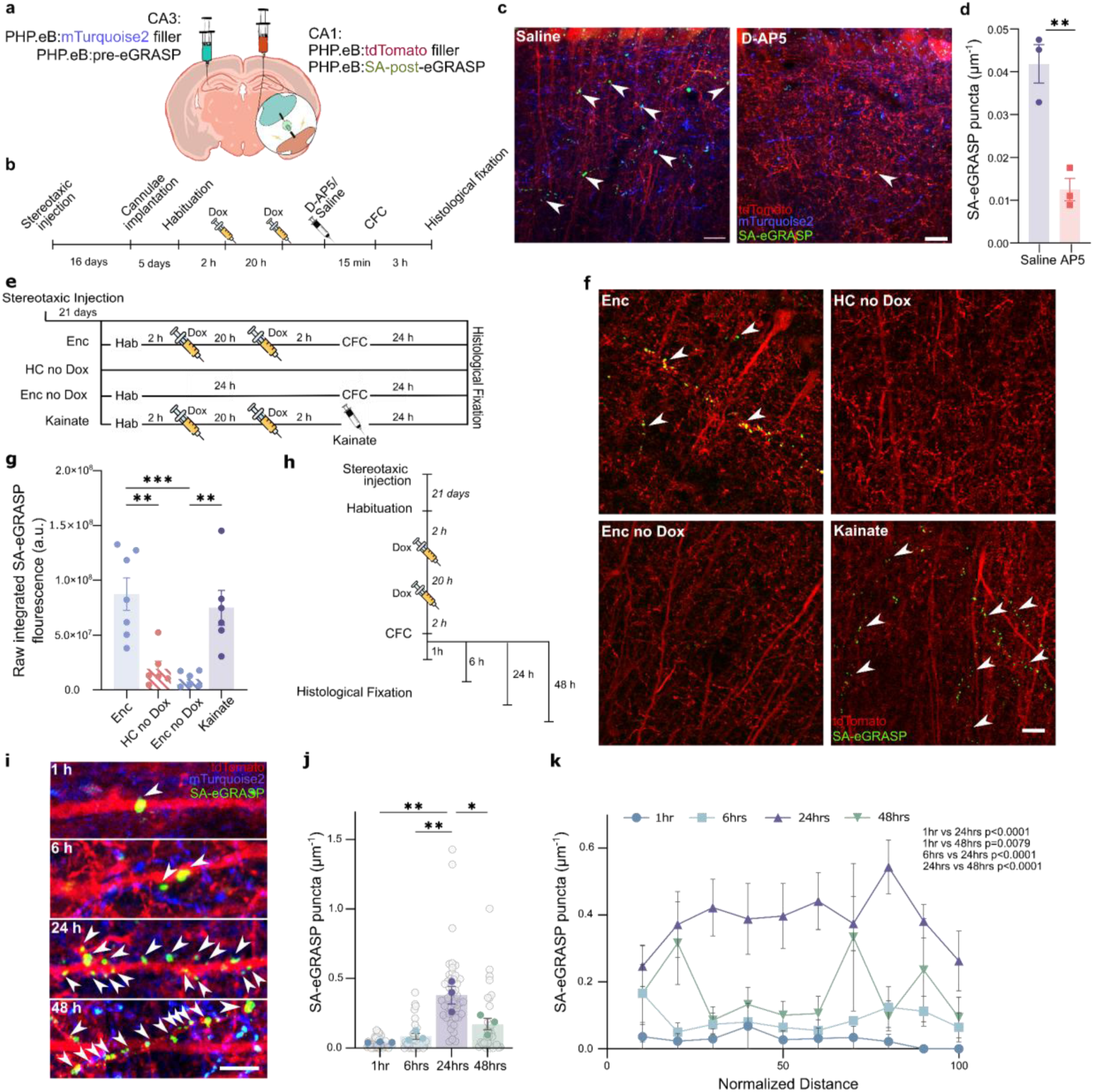
SA-eGRASP expression depends on activity. **a,b.** Description of the gene constructs and of the AAV stereotaxic injection sites (a); experimental timeline for local D-AP5 or saline injection (b). **c.** Representative confocal images (30 z-planes) of SA-eGRASP expression (arrows) following local D-AP5 or saline injection. Scale bar 10 μm. **d**. Density of SA-eGRASP+ puncta on tdTomato+ dendrites (unpaired, two-tailed t-test, *t*_4_ = 5.608, *P* = 0.0050; N_Saline_ = 3, and N_AP5_ = 3 mice per group). **e.** Experimental timeline for Encoding with Dox (Enc), Homecage without Dox (HC no Dox), Encoding without Dox (Enc no Dox), and Kainate control mouse groups. **f.** Representative confocal images (30 z-planes) of SA-eGRASP expression. Scale bar 10 μm. **g.** Raw integrated density of SA-eGRASP fluorescence following Enc, HC no dox, Enc no Dox, and Kainate controls (ANOVA-1, *F*_3,21_ = 10.92, *P* = 0.0002; Tukey’s multiple comparisons test, Enc no dox vs Enc *P* = 0.0007, Enc no dox vs Kainate *P* = 0.0053, Enc vs HC no dox *P* = 0.0025, HC no dox vs Kainate *P* = 0.0186; N_Enc_= 7, N_Hc no Dox_ = 6, N_Enc no Dox_ = 6, N_Kainate_ = 6 sections; 3 mice per group). **h.** Experimental timeline for investigating the time course of SA-eGRASP expression. **i.** Representative confocal images (10 z-planes) of SA-eGRASP expression 1, 6, 24, and 48 h after CFC. Scale bar, 5 μm. **j**. SA-eGRASP+ puncta density 1, 6, 24, and 48 h after CFC (full circles: mean value per mouse; empty circles: values per individual dendritic trace analyzed; ANOVA-1, *F*_3,8_ = 14.38, *P* = 0.0014; Tukey’s multiple comparisons test, *P*_1 h vs 24 h_ = 0.0014, *P*_6 h vs 24 h_ = 0.0033, *P*_24 h vs 48 h_ = 0.0255; N = 2 sections per mouse; N = 3 mice for all experimental groups). **k.** Density of SA-eGRASP+ puncta across the stratum radiatum at different time points (0 is soma position and 100 is end of stratum radiatum; ANOVA-2, time, *F*_3,114_ = 34.80, *P* < 0.0001; Tukey’s multiple comparisons test, *P*_1 h vs 24 h_ < 0.0001, *P_1_* _h vs 48 h_ = 0.0079, *P*_6 h vs 24 h_ < 0.0001, *P*_24 h vs 48 h_ < 0.0001; N = 2 sections per mouse; N = 3 mice for each experimental group).

**Extended Data Figure 2.**
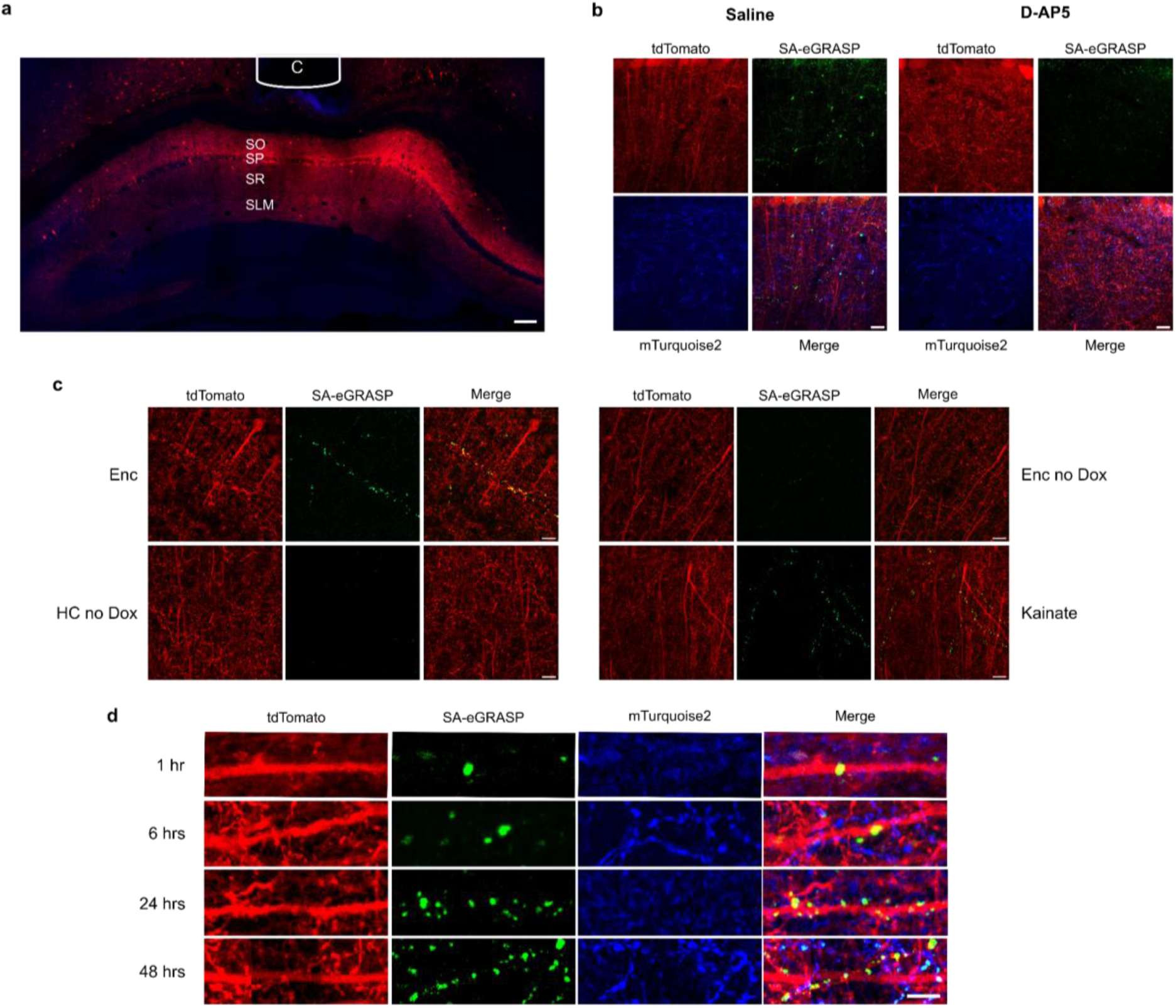
Confocal images of SA-eGRASP *ex vivo* expression controls. **a**. Representative low-magnification confocal image indicating cannula (C) placement above the hippocampal CA1 region (SO, Stratum Oriens; SP, Stratum Pyramidale; SR, Stratum Radiatum; SLM, Stratum Lacunosum Moleculare). Scale bar, 100 μm. **b.** Representative confocal images of SA-eGRASP expression following local saline or D-AP5 injection. Scale bar, 10 μm. **c.** Representative images of SA-eGRASP expression in Enc, HC-no Dox control, Enc-no Dox, and Kainate controls. Scale bar, 10 μm. **d**. Representative confocal images of SA-eGRASP expression 1, 6, 24, and 48 h after CFC. Scale bar, 5 μm.

**Extended Data Figure 3.**
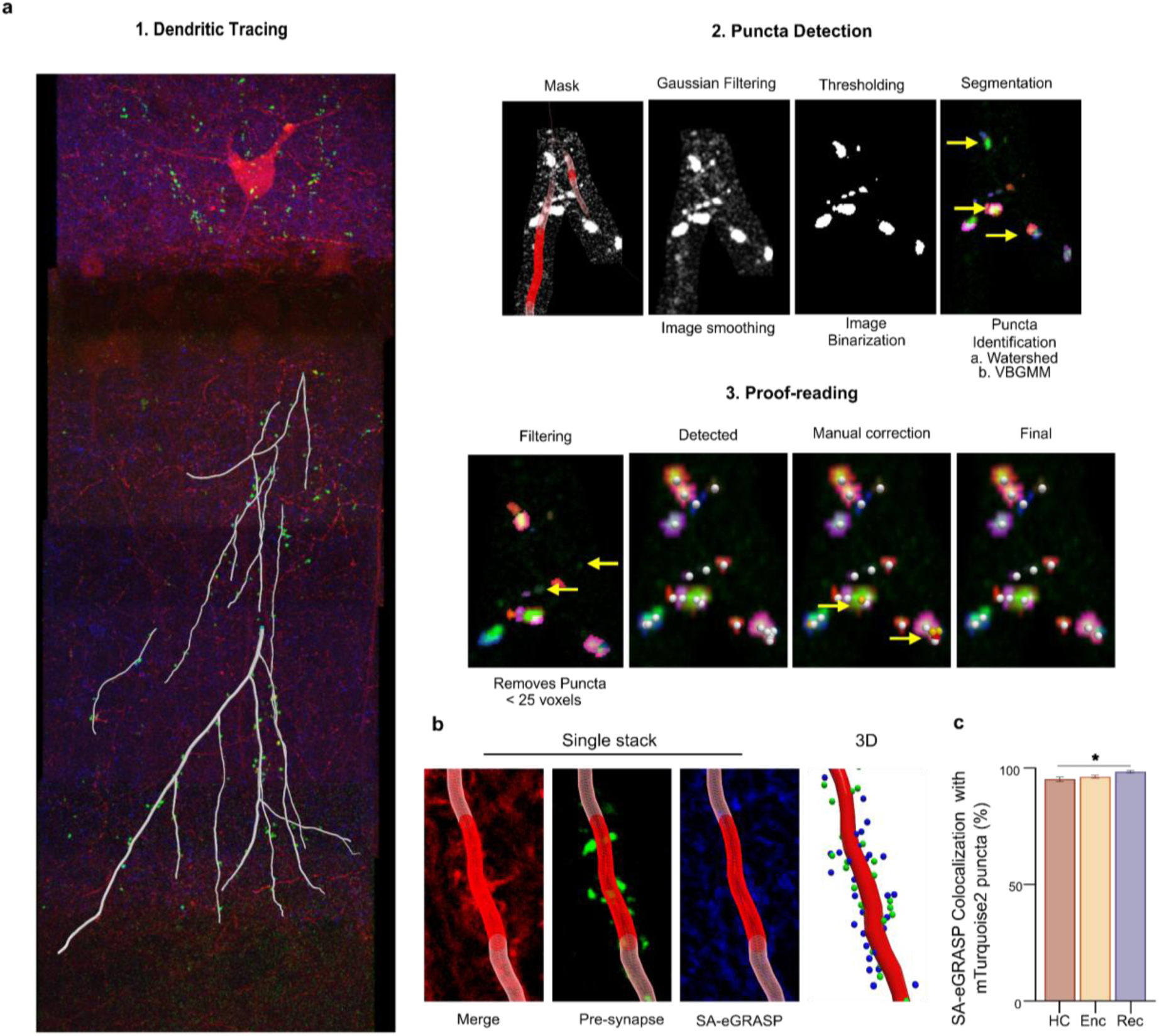
Pipeline for Semi-Automated detection of SA-eGRASP puncta. **a.** Detection of SA-eGRASP+ puncta. **a.1**) Representative confocal microscopy image of the CA1 region with superimposed tdTomato+ CA1 dendrites traces, reconstructed using Neutube, of dendrites expressing SA-eGRASP signal reconstructed using Neutube. **a.2**) Stepwise process of puncta detection using custom MATLAB code: the dendritic trace is used to create a mask delimiting the voxels analyzed, which are first smoothened through Gaussian filtering. Voxels with intensities greater than I_min_ are then binarized. Binarized voxels larger than V_max_ (yellow arrow) are segmented by watershed and/or mean-shift-regulated Variational Bayesian Gaussian Mixture Models. **a.3**) Identified puncta with a voxel size smaller than V_min_ are filtered (yellow arrows). Individual puncta identified after segmentation and filtering are marked in different colors, with white spheres marking their centers. Over-segmented puncta may be manually corrected (yellow arrows). **b**. Representative confocal microscopy images showing a single-stack image of dendrite, and corresponding masked blue signal (pre-synaptic mTurquoise2) and green signal (SA eGRASP). 3-D visualization (right) of detected puncta in blue and green channels, showing their colocalization. **c.** Percentage of detected green (i.e., SA-eGRASP+) puncta that show colocalization (distance between centroids (d) < 2 μm) with detected blue (i.e., mTurquoise2^+^) puncta. (N_HC_ = 7, N_Enc_ = 6, N_Rec_ = 6 mice).

To check the dependence on NMDAR activation of SA-post-eGRASP expression also *in vivo*, we injected D-AP5 into the dCA1 prior to CFC (Fig. 2b,c; Extended Data Fig. 2a). This significantly reduced the density of SA-eGRASP+ puncta compared to saline-injected controls (Fig. 2c,d; Extended Data Fig. 2b), indicating that SA-eGRASP expression depends on NMDAR activation. When we omitted doxycycline, the SA-eGRASP signal was negligible even after CFC (Fig. 2e,g; Extended Data Fig. 2c), demonstrating virtually zero leakage of the system. Finally, as a positive control for activity-dependent induction, we systemically administered kainic acid to increase neuronal activity. This resulted in robust SA-eGRASP labeling (Fig. 2e-g; Extended Data Fig. 2c)^22^.

To assess the temporal dynamics of SA-eGRASP expression, we performed histological fixation at different time points following CFC (Fig. 2h). We then quantified SA-eGRASP+ puncta by confocal microscopy using a custom, semi-automated analysis pipeline (Extended Data Fig. 3). SA-eGRASP+ puncta were evident by intrinsic fluorescence and were distributed along tdTomato+ dCA1 dendrites, colocalizing with mTurquoise2+ commissural dCA3 axons (Fig. 2i; Extended Data Fig. 2d). SA-eGRASP+ puncta density peaked at 24 h after CFC throughout the stratum radiatum (SR) and started to decline 48 h post-CFC (Fig. 2j,k).

These results show that, consistent with our findings in primary hippocampal cultures, SA-eGRASP labels robustly and selectively potentiated synapses *in vivo* in an NMDAR- and doxycycline-dependent manner.

### Encoding and recall of a contextual fear memory increase the number, size, and clustering of SA-eGRASP+ synapses across the dCA1 Stratum Radiatum

To map the spatial distribution of potentiated dCA3 to dCA1 synapses along Stratum Radiatum (SR) dendrites associated with distinct stages of memory formation, we subjected three groups of mice to different behavioral paradigms and labelling schedules. The Homecage (HC) group served as a control to measure baseline synaptic potentiation. The Encoding (Enc) group captured synapses potentiated during the encoding phase of learning - when the contextual fear memory is acquired, and the Recall (Rec) group captured synapses potentiated during memory retrieval upon re-exposure to the conditioning context 24 h later (Fig. 3a,b; Extended Data Fig. 4).

Our custom, semi-automated image analysis pipeline (Extended Data Fig. 3a,b) enabled us to reconstruct the spatial distribution of potentiated synapses by providing the spatial coordinates and volume of each SA-eGRASP+ punctum along the apical dendritic tree. We found a significant increase in SA-eGRASP+ puncta density following Enc compared to HC (Fig. 3c). The interindividual variability of puncta density was higher among Enc and Rec mice relative to HC (Fig. 3d). The percentage of SA-eGRASP+ puncta normalized to mTurquoise2+ dCA3 boutons was also higher in Enc mice than in HC and Rec (Fig. 3e) across the whole SR (Fig. 3f). The density of mTurquoise2+ dCA3 presynaptic boutons did not differ between groups (Fig. 3g,h). No significant correlation was observed between SA-eGRASP+ puncta density and freezing behavior duration (Extended Data Fig. 5).

These findings reveal that memory formation is accompanied by stage-specific patterns of synaptic potentiation within the dCA3–dCA1 circuit. Learning-induced plasticity is spatially organized along the dendritic arbor of individual dCA1 neurons, highlighting how distinct phases of memory formation reshape hippocampal connectivity at the level of specific synapses.

GRASP+ puncta size correlates strongly with synaptic morphology and functional connectivity^23^. We therefore used SA-eGRASP+ puncta volume as a proxy for synaptic potentiation^24^. Puncta volumes were larger in Enc and Rec relative to HC groups, with Rec values comparable to Enc (Fig. 3i). This increase was accompanied by a distal–proximal gradient along the SR, with larger puncta located at greater distances from the soma in Enc and Rec mice (Extended Data Fig. 6). To determine whether SA-eGRASP expression detected after recall might also include a quota of puncta that had actually been generated after the CFC encoding phase, a group of mice received identical doxycycline administration to Rec but was not re-exposed to the context (Recall-Blank, RecBl). Surprisingly, RecBl animals showed reduced SA-eGRASP+ puncta density compared to HC (Extended Data Fig. 7a-f). Moreover, exposure to handling alone (Handl) resulted in SA-eGRASP levels comparable to HC (Extended Data Fig. 7a-f). Presynaptic bouton density remained unchanged across all groups (Extended Data Fig. 7g,h).

To examine whether the spatial distribution of SA-eGRASP+ puncta changes after the CFC experience, we analyzed differences in the nearest-neighbor distances (NNDs), *i.e.* the pathlength distance (d) between two adjacent SA-eGRASP+ puncta within the same dendritic segment (Fig. 4a), among the different groups. NNDs in the Enc group were significantly smaller than in the HC group, with a similar trend in the Rec group (Fig. 4b). SA-eGRASP+ puncta in HC, Enc, and Rec groups were more clustered than corresponding control distributions generated via data shuffling (Fig. 4c). We then defined a cluster as a group of three or more SA-eGRASP+ puncta whose NNDs were below the median value observed in the HC group (median d_HC_ = 1.16 μm). Dendritic segments in the Enc and Rec groups showed a trend towards higher cluster density compared to HC (Fig. 4d), with a significantly higher percentage of SA-eGRASP+ puncta belonging to a cluster in Enc as compared to HC (Fig. 4e). The Rec group showed a similar trend (Fig. 4e). While we observed a significant difference between groups, we found no significant difference in volume between clustered and isolated SA-eGRASP+ puncta within groups (Fig. 4f). The number of SA-eGRASP+ puncta composing each cluster was significantly larger in Enc vs. HC, indicating larger cluster size (Fig. 4g).

**Figure 3.**
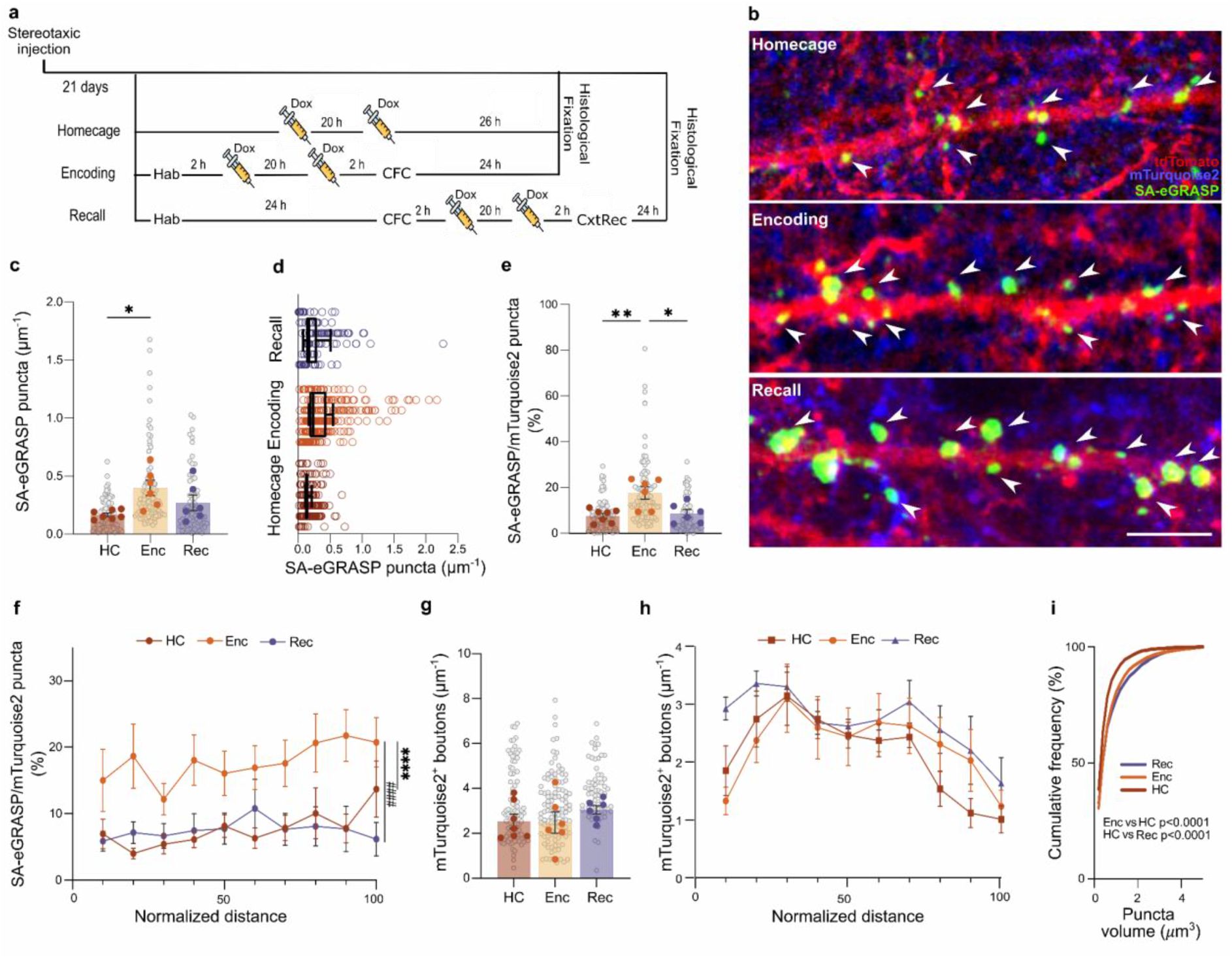
SA-eGRASP labels synapses potentiated upon learning. **a,b**. Experimental timeline (a) and representative confocal microscopy images (10 z-planes) showing CA1 dendrites (red), CA3 axons (blue), and SA-eGRASP puncta (green) in the dCA1 of mice after home cage housing (HC), contextual fear conditioning encoding (Enc), or contextual fear conditioning recall (Rec). Habituation, Hab; Contextual fear conditining, CFC; Contextual recall, CxtRec. Scale bar, 5 μm. **c.** Density of SA-eGRASP+ puncta (ANOVA-1, *F*_2,16_ = 4.918, *P* = 0.0216; Tukey’s multiple comparisons tests, *P*_HC vs Enc_ = 0.0166). **d**. SA-eGRASP density variability within branches across HC, Enc, and Rec groups. Each point represents a dendrite section within an individual mouse (row). Box-and-whisker plots summarize the distribution of within-mouse branch-density variability across groups, showing the median, IQR, and minimum-to-maximum range. Relative to HC, Enc and Rec exhibited ∼2.4-fold and ∼2.0-fold greater variability, respectively (likelihood-ratio test, χ²_2_ = 147.0, *P* < 0.0001). **e**. SA-eGRASP+ puncta increase after Enc (ANOVA-1, *F*_2,16_ = 8.1, *P* = 0.0035; Tukey’s multiple comparisons test, *P*_HC vs Enc_ = 0.0047, *P*_Enc vs Rec_ = 0.0131). **f**. Percentage of SA-eGRASP+ puncta along the SR (0 and 100% correspond to the borders with SP and SLM, respectively; ANOVA-2, experimental group, *F*_2,160_ = 37.98, *P* < 0.0001; Tukey’s multiple comparisons test, *P*_Enc vs HC_ < 0.0001, *P*_Enc vs Rec_ < 0.0001) **g,** Density of detected mTurquoise2+ CA3 boutons in the dCA1 SR (ANOVA-1, *F*_2,16_ = 0.7758, *P* = 0.4769). **h**. Distribution of mTurquoise2+ CA3 boutons along the CA1 SR (ANOVA-2, experimental group, *F*_2,159_ = 5.996, *P* = 0.0031; Tukey’s multiple comparisons test at 10% distance, Enc vs Rec ^##^*P* = 0.0119). **i**. Cumulative distribution of detected SA-eGRASP+ puncta volumes. Only volumes below 5 μm^3^ are included. (Kolmogorov-Smirnov test, Enc vs HC *D* = 0.1305, *P* < 0.0001, HC vs Rec *D* = 0.1405, *P* < 0.0001; N_HC_ = 1971, N_Enc_ = 5847, N_Rec_ = 1768). For panels **c**, **e**, and **g**, grey circles represent individual dendrite sections (N_HC_ = 94 traces from 7 mice, N_Enc_ = 107 traces from 6 mice, N_Rec_ = 66 traces from 6 mice).

**Extended Data Figure 4.**
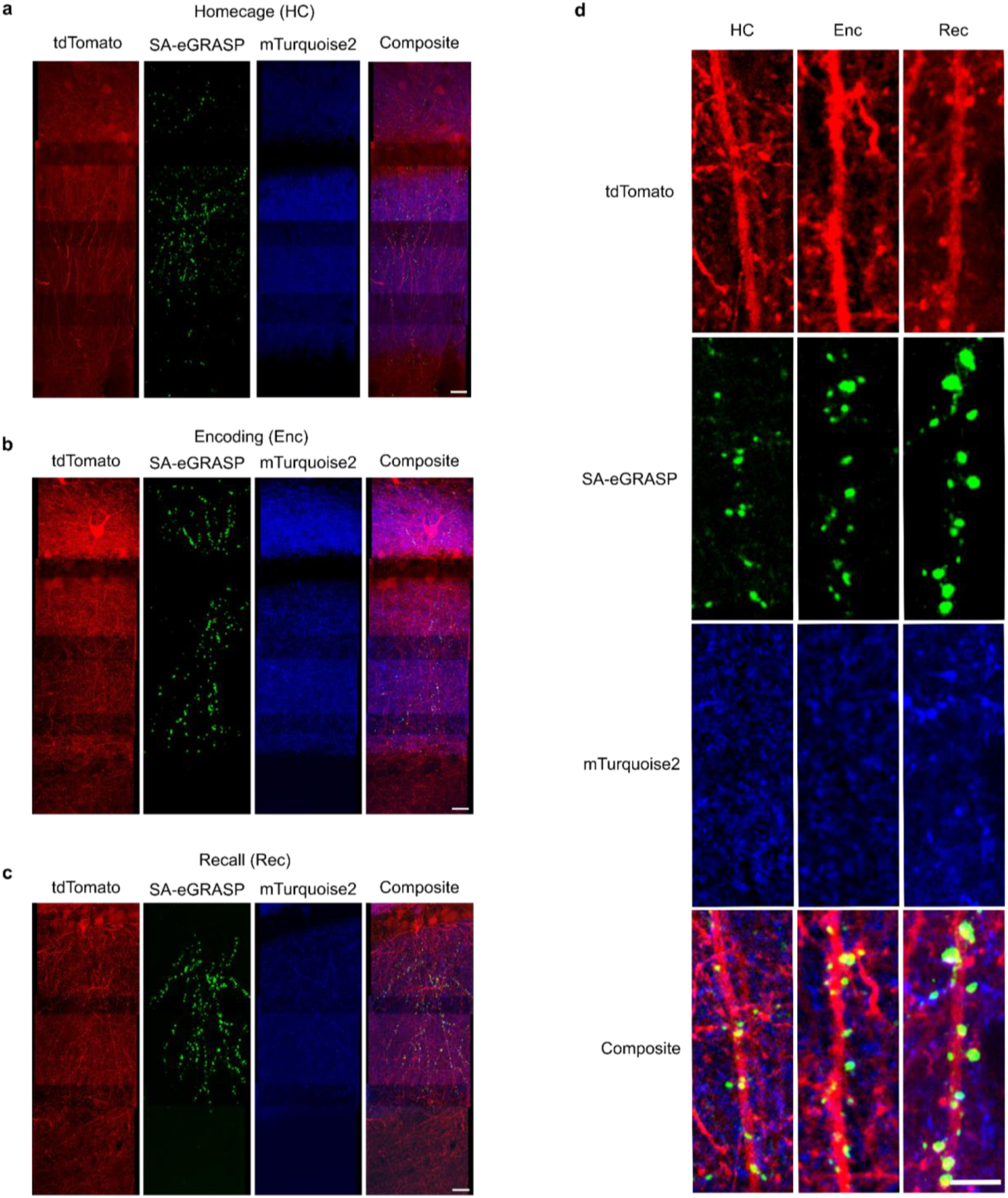
Confocal microscopy imaging of SA-eGRASP expression along the stratum radiatum. Representative images showing CA1 dendrites (red), CA3 axons (blue), and SA-eGRASP (green) of mice (**a**) kept in their home cage (HC), (**b**) subjected to the association phase of the contextual fear conditioning paradigm (encoding, Enc), (**c**) re-exposed to the conditioning chamber (recall, Rec). Scale bar, 20 μm. **d.** Higher-magnification images showing SA-eGRASP expression in HC, Enc, and Rec. Scale bar, 5 μm.

**Extended Data Figure 5.**
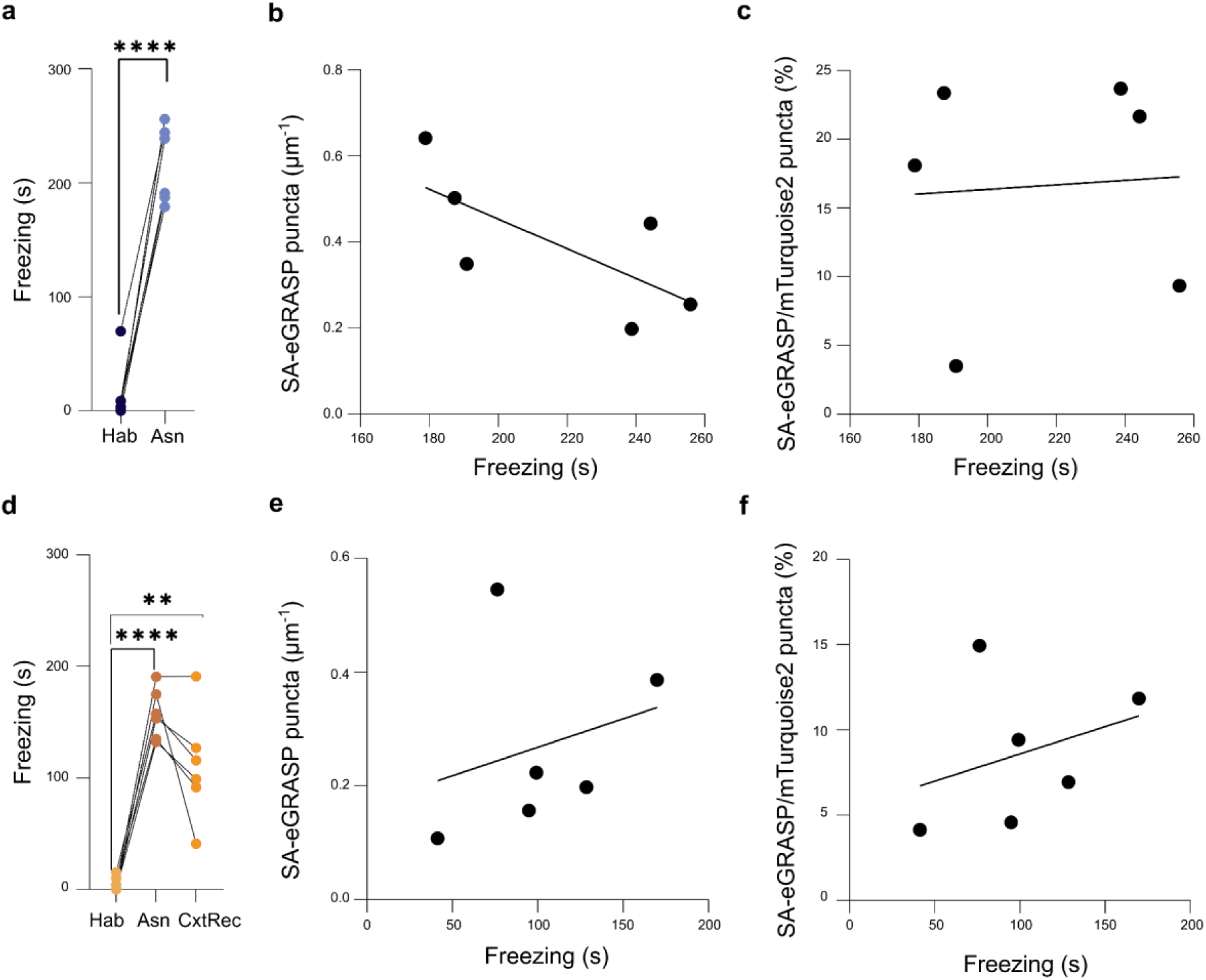
SA-eGRASP expression in relation to behavioral output. **a**. Time freezing during habituation (Hab) and context association (Asn) for the Enc experimental group (paired t-test, *P*_Hab vs Asn_ = 0.0031). **b**. Relationship between time freezing and SA-eGRASP density for the Enc group (simple linear regression, *R²* = 0.5048, *F*_1,4_ = 4.078, *P* = 0.1136; N_Enc_ = 6). **c**. Relationship between time freezing and SA-eGRASP percentage for the Enc group (simple linear regression, *R²* = 0.004475, *F*_1,4_ = 0.01798, *P* = 0.8998; N_Enc_ = 6). **d**. Time freezing during habituation (Hab), context association (Asn), and contextual recall (CxtRec) for the Rec experimental group (repeated-measures ANOVA-2, *F*_1.398, 6.992_ = 43.33, *P* = 0.0002; Tukey’s multiple comparisons test, *P*_Hab vs Asn_< 0.0001, *P*_Hab vs. Rec_ = 0.0073; N = 6 per group). **e.** Relationship between time freezing and SA-eGRASP density for the Rec experimental group (simple linear regression, *R²* = 0.07145, *F*_1,4_ =0.3078, *P* = 0.6086; N_Rec_ = 6). **f.** Relationship between time freezing and SA-eGRASP percentage for the Rec experimental group (simple linear regression, *R²* = 0.1100, *F*_1,4_ = 0.4945, *P* = 0.5207; N_Rec_ = 6).

**Extended Data Figure 6.**
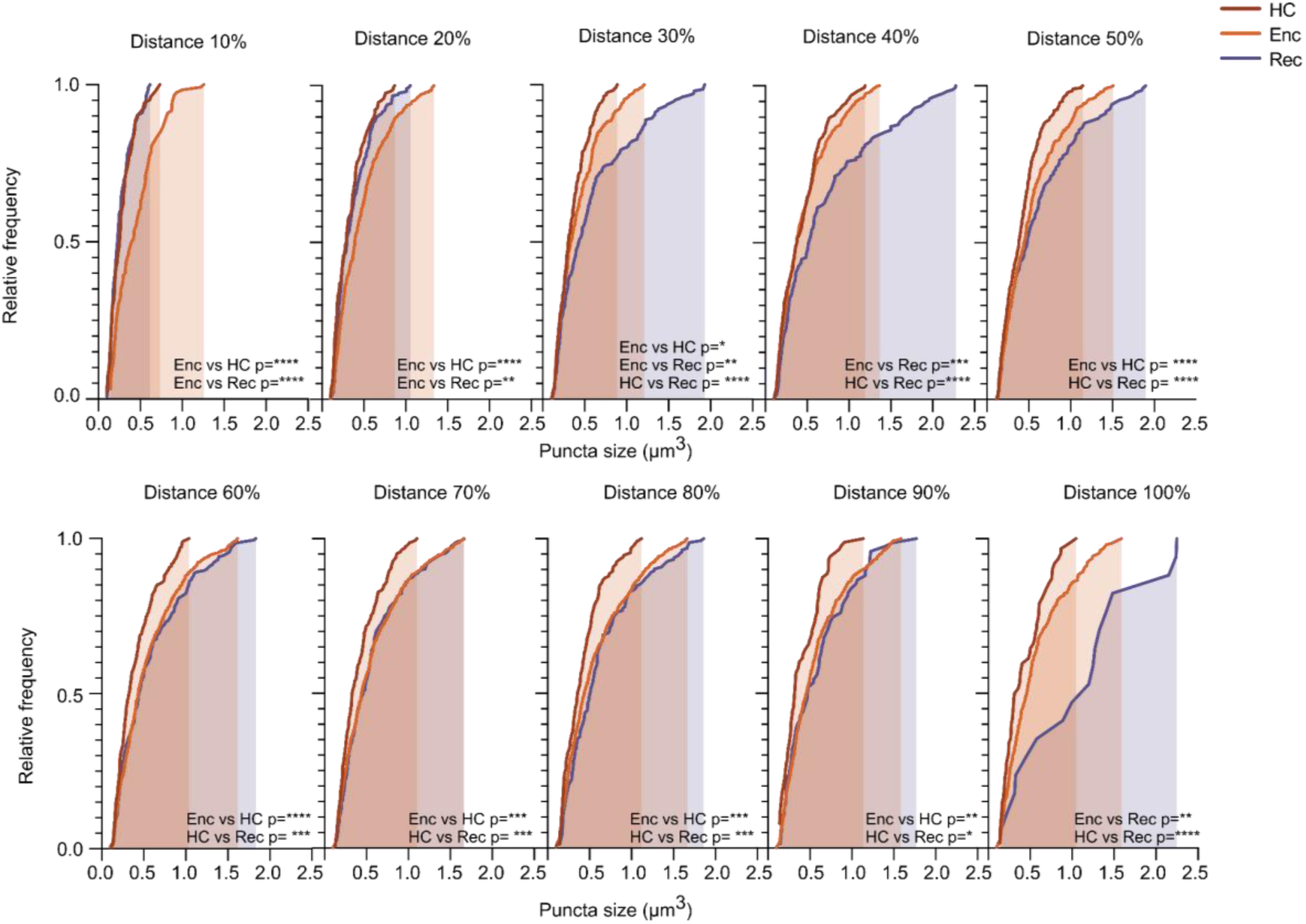
Distribution of SA-eGRASP^+^ puncta volume along the Stratum Radiatum. Cumulative distributions of the volumes of detected SA-eGRASP+ puncta as a function of their position along the Stratum Radiatum, calculated at 10% intervals of the normalized distance between the borders with Stratum Pyramidale (0% distance) and Stratum Lacunosum Moleculare (100% distance). Only volumes below 5 μm^3^ are included. (Kolmogorov-Smirnov test, distance 10%, Enc vs HC, *D* = 0.3436, *P* < 0.0001, Enc vs Rec, d = 0.3440, *P* < 0.0001; distance 20%, Enc vs HC, *D* = 0.2203, *P* < 0.0001, Enc vs Rec, *D* = 0.1582, *P* = 0.0014; distance 30%, Enc vs HC, *D* = 0.1186, *P* = 0.0377, Enc vs Rec, *D* = 0.1609, *P* = 0.0018, HC vs Rec, *D* = 0.2413, *P* < 0.0001; distance 40%, Enc vs Rec, *D* = 0.1883, *P* = 0.0002, HC vs Rec, *D* = 0.2386, *P* < 0.0001; distance 50%, Enc vs HC, *D* = 0.1396, *P* = 0.0006, HC vs Rec, *D* = 0.2183, *P* = 0.0003; distance 60%, Enc vs HC, *D* = 0.1771, *P* < 0.0001, HC vs Rec, *D* = 0.1880, *P* = 0.0005; distance 70%, Enc vs HC, *D* = 0.1603, *P* = 0.0002, HC vs Rec, *D* = 0.1883, *P* = 0.0007; distance 80%, Enc vs HC, *D* = 0.1938, *P* = 0.0004, HC vs Rec, *D* = 0.2377, *P* = 0.0005; distance 90%, Enc vs HC, *D* = 0.2167, *P* = 0.0014, HC vs Rec, *D* = 0.2429, *P* = 0.0168; distance 100%, Enc vs Rec, *D* = 0.4605, *P* = 0.0021, HC vs Rec, *D* = 0.6148, *P* < 0.0001; number of puncta analyzed for each distance: 10%, N_HC_ = 104, N_Enc_ = 122, N_Rec_ = 113; 20%, N_HC_ = 168, N_Enc_ = 463, N_Rec_ = 213; 30%, N_HC_ = 220, N_Enc_ = 393, N_Rec_ = 207; 40%, N_HC_ = 234, N_Enc_ = 624, N_Rec_ = 170; 50%, N_HC_ = 305, N_Enc_ = 653, N_Rec_ = 135; 60%, N_HC_ = 270, N_Enc_ = 783, N_Rec_ = 204; 70%, N_HC_ = 233, N_Enc_ = 877, N_Rec_ = 215; 80%, N_HC_ = 133, N_Enc_ = 787, N_Rec_ = 164; 90%, N_HC_ = 88, N_Enc_ = 626, N_Rec_ = 75; 100%, N_HC_ = 62, N_Enc_= 327, N_Rec_= 17; N_HC_ = 7, N_Enc_ = 6, N_Rec_ = 6 mice).

**Extended data figure 7.**
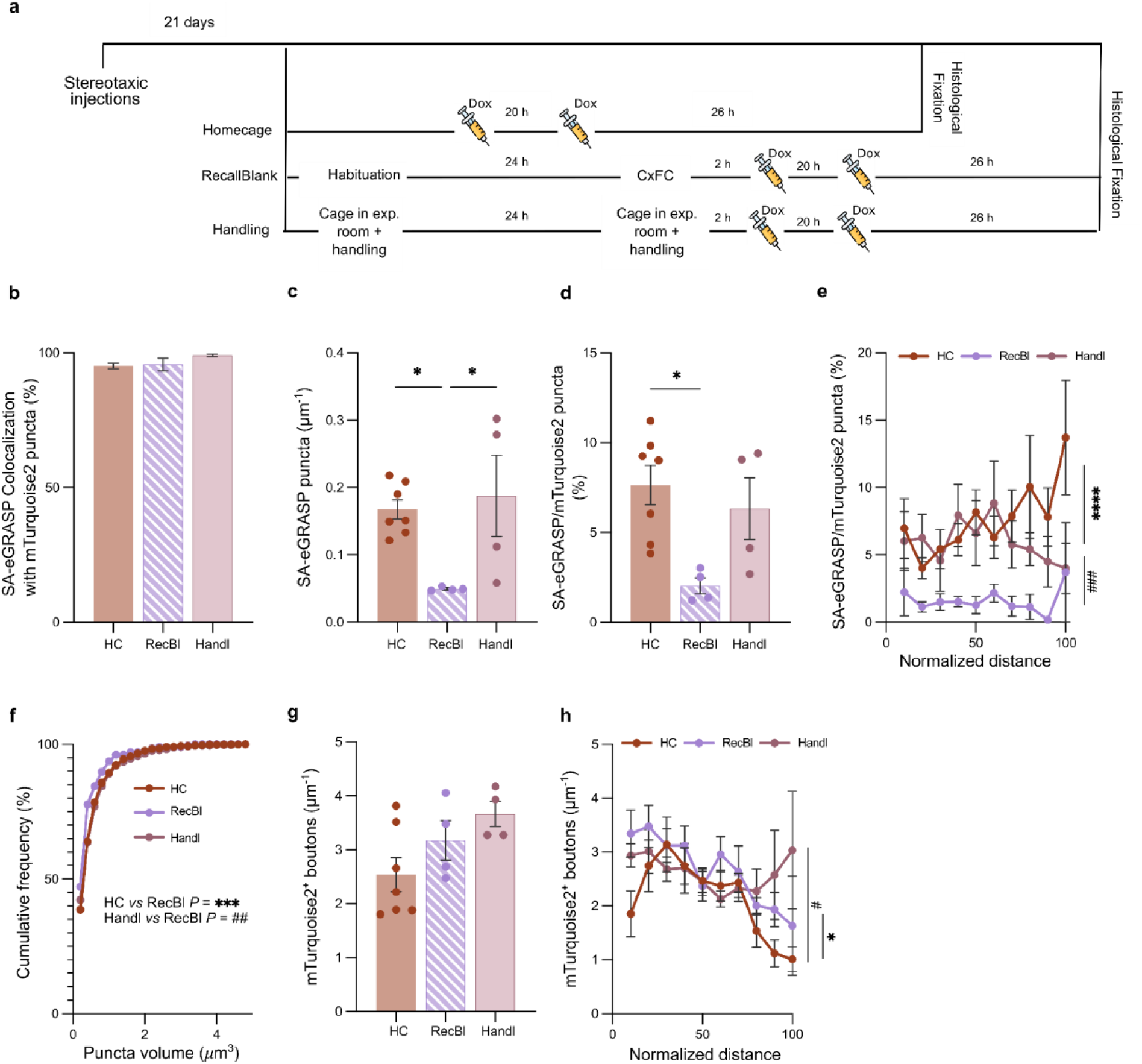
Reduction in expression at Recall blank is not due to a habituation to handling. **a.** Schematic of experimental timelines. **b**. Percentage of detected green (i.e., SA-eGRASP+) puncta showing colocalization (distance between centroids (d) < 2 μm) with detected blue (i.e., mTurquoise2+) puncta. **c.** Density of detected SA-eGRASP+ puncta (ANOVA-1, *F*_2,12_ = 5.395, *P* = 0.0213; Tukey’s post-hoc test, *P*_HC vs RecBl_ = 0.0365, *P*_Handl vs RecBl_ = 0.0299; N_HC_ = 7, N_RecBl_ = 4, N_Handl_ = 4 mice). **d**. Percentage of SA-eGRASP+ puncta detected (ANOVA-1, *F*_2,12_, *P* = 0.0191; Tukey’s post-hoc test, *P*_HC vs RecBl_ = 0.0157; N_HC_ = 7, N_RecBl_ = 4, N_Handl_ = 4 mice). **e**. Distribution of the percentage of SA-eGRASP+ puncta along the stratum radiatum (ANOVA-2, experimental group, *F*_2,115_=16.85 *P* < 0.0001; Tukey’s post-hoc test, P_HC vs RecBl_ < 0.0001, P_Handl vs RecBl_ = 0.0009; N_HC_ = 7, N_RecBl_ = 4, N_Handl_ = 4 mice). **f**. Cumulative distribution of detected SA-eGRASP+ puncta volumes across the CA1 stratum radiatum. Only volumes below 5 μm^3^ are included (Kolmogorov-Smirnov test, HC vs RecBl,d = 0.1468, *P* = 0.0006, Handl vs RecBl, *D* = 0.1462, *P* = 0.0016; N_HC_=1971, N_RecBl_ = 206, N_Handl_ = 894 puncta). **g**. Density of detected mTurquoise2+ CA3 presynaptic boutons in the CA1 stratum radiatum normalized to the dendritic trace length (ANOVA-1, *F*_2,12_ = 3.183, *P* = n.s.; N_HC_ = 7, N_RecBl_ = 4, N_Handl_ = 4 mice). **h**. Distribution along the stratum radiatum of detected mTurquoise2+ CA3 presynaptic boutons normalized to the dendritic trace length (ANOVA-2, normalized distance, *F*_9,115_ = 3.413, *P* = 0.0009; experimental group, *F*_2,115_ = 5.796, *P* = 0.0040; Tukey’s post-hoc test, distance 10%, *P*_HC vs RecBl_ = 0.0196, distance 90%, *P*_HC vs Handl_ = 0.0441, distance 100%, *P*_HC vs Handl_ = 0.0038; N_HC_ = 7, N_RecBl_ = 4, N_Handl_ = 4 mice).

**Figure 4.**
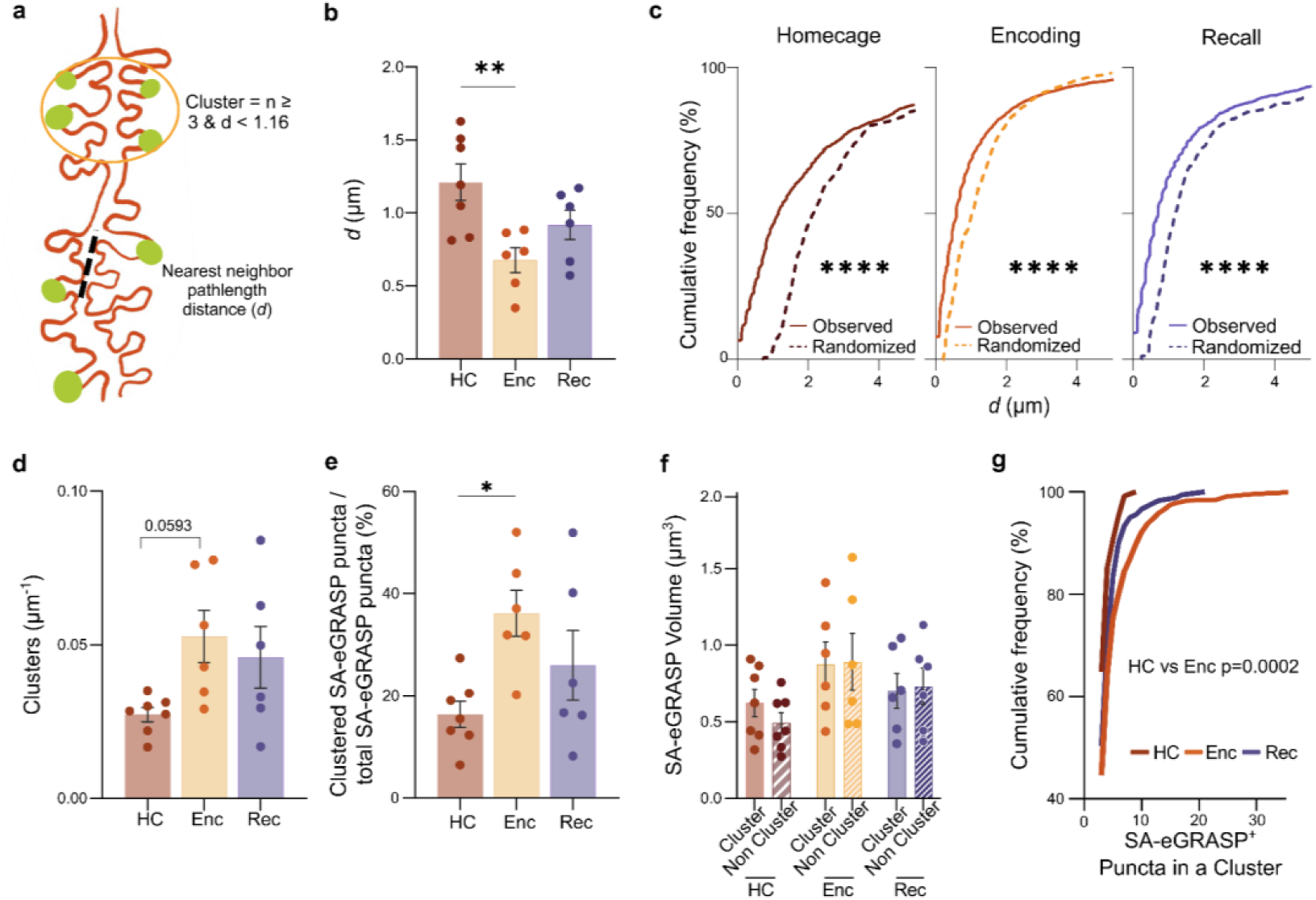
Clustering of SA-eGRASP+ puncta following learning. **a**. Representation of the dendritic pathlength between two adjacent SA-eGRASP+ puncta (Nearest-Neighbor pathlength Distance, d), with parameters for defining a synapse cluster. **b**. Nearest-Neighbor pathlength Distance, (d) median values for HC, Enc and Rec groups; each data point is the median value for one mouse (ANOVA-1, *F*_2,16_ = 6.344, *P* = 0.0094; N_Enc_ = 6, N_HC_ = 7, N_Rec_ = 6 mice). **c**. Cumulative distribution of experimental versus shuffled *d* values (Kolmogorov-Smirnov test, HC, real vs random, *D* = 0.4385, *P* < 0.0001, Enc, real vs random, *D* = 0.2548, *P* < 0.0001; Rec, real vs random, *D* = 0.1115, *P* < 0.0001). **d**. Density of SA-eGRASP+ puncta normalized to dendritic pathlength (ANOVA-1, *F*_2,16_ = 3.395, *P* = 0.0590; Tukey’s post-hoc test, *P*_HC vs Enc_ = 0.0593). **e**. Percentage of SA-GRASP+ puncta belonging to a cluster (ANOVA-1, *F*_2,16_ = 4.487, *P* = 0.0284; Tukey’s post-hoc test, *P*_Enc vs. HC_ = 0.0221). **f.** Comparison of volumes of SA-eGRASP+ puncta belonging or not to clusters (ANOVA-2, experimental group, *F*_2,32_ = 3.662, *P* = 0.0370). **g.** Cumulative distribution of the number of SA-eGRASP+ forming a cluster (Kolmogorov-Smirnov test, HC vs Enc, *D* = 0.2080, *P* = 0.0002; N_HC_ = 128 clusters from 7 mice, N_Enc_ = 592 clusters from 6 mice, N_Rec_ = 181 clusters from 6 mice).

These findings show that clustering of SA-eGRASP+ puncta is associated with learning-induced synaptic potentiation of dCA3–dCA1.

The high signal-to-noise ratio of the SA-eGRASP signal prompted us to investigate the dynamics of CA3-to-CA1 SA-eGRASP+ synapses over extended temporal windows using longitudinal two-photon time-lapse imaging in the SO of dCA1. To visualize individual SA-eGRASP+ spines, we transduced the right dCA1 neurons with three AAVs encoding Cre recombinase under the CaMKII promoter, Cre-dependent tdTomato, rtTA under the CAG promoter, and SA-post-eGRASP. Contralateral CA3 neurons were transduced with AAVs encoding rtTA under the hSyn promoter and pre-eGRASP (Fig. 5a). To induce synaptic potentiation, we exposed mice to an enriched environment (EE) for 16 h (Fig. 5b), and doxycycline was administered 24 h before and immediately prior to EE exposure. To check the leakage of this system, we included two control groups: mice exposed to EE injected with vehicle only, and mice administered doxycycline without exposure to EE (Fig. 5b). As these groups exhibited indistinguishable structural dynamics (Extended Data Fig. 8a,b), we pooled them into a single control group for subsequent analyses.

**Figure 5.**
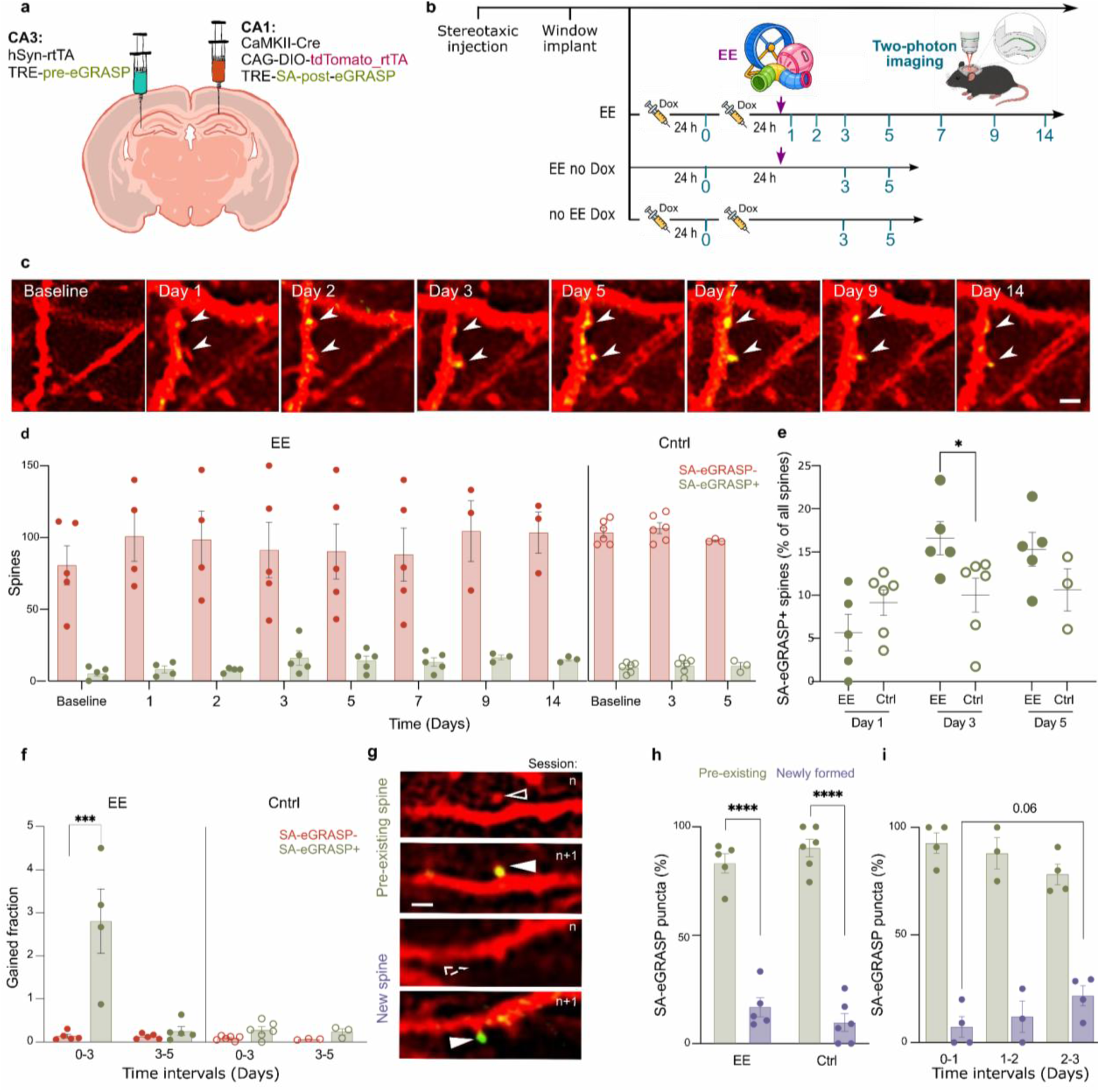
Longitudinal tracking of SA-eGRASP expression at individual dendritic spines. **a,b.** Description of the gene constructs and of the AAV stereotaxic injection sites (a); experimental timeline for tracking a sparse subset of dCA1 pyramidal neurons and SA-eGRASP puncta on their dendritic spines by two-photon optical imaging (b). **c.** 2P time series of dCA1 pyramidal neuron dendrites expressing tdTomato (red) and SA-eGRASP puncta (green) over 14 days (single z-plane). Scale bar, 2 μm. **d**. Number of eGRASP-negative (red) and eGRASP-positive (green) spines per mouse in the EE and Control group over time. **e.** Percentage of SA-eGRASP+ in EE (solid circles) and Control (empty circles) groups at different time points (ANOVA-1, *F*_5,24_ = 4.380*, P* = 0.0056; Fisher’s LSD post-hoc test, *P*_Day1_Cntrl vs Day1_EE_ = 0.2014, *P*_Day3_Cntrl vs Day3_EE_ = 0.0200, *P*_Day5_Cntrl vs Day5_EE_ = 0.1529; each data point is one mouse). **f**. Fraction of newly formed SA-eGRASP+ (green) and SA-eGRASP-(red) spines per mouse for both EE and Control groups (ANOVA-1, *F*_7,29_ = 14.27, *P* < 0.0001; Holm–Šídák post-hoc test, *P*_Int. 0-3, EE SA-eGRASP-vs EE SA-eGRASP+_ < 0.0001; each data point is one mouse). **g**. Representative 2P images (single z-plane) of SA-eGRASP puncta (green) formed on either new or pre-existing spines of dCA1 neurons labeled with tdTomato (red). Scale bar, 2 μm. **h**. Percentage of SA-eGRASP+ puncta per mouse appearing on new and pre-existing spines, all time points pooled (ANOVA-1, *F*_3,18_ = 102.5, *P* < 0.0001; Holm–Šídák post-hoc test, *P*_Pre existing_EE vs Newly formed_EE_ < 0.0001, *P*_Pre existing_Cntrl vs Newly formed_ Cntrl_ < 0.0001; each data point represents one mouse). **i.** SA-eGRASP puncta formed on newly generated and on pre-existing spines within the EE group across days (ANOVA-1, *F*_5,16_ = 55.54, *P* < 0.0001; Fisher’s LSD post-hoc test, *P*_Newly formed_ int 0-1 vs. Newly formed_int 2-3_ = 0.0646; each data point represents one mouse).

**Extended Data Figure 8.**
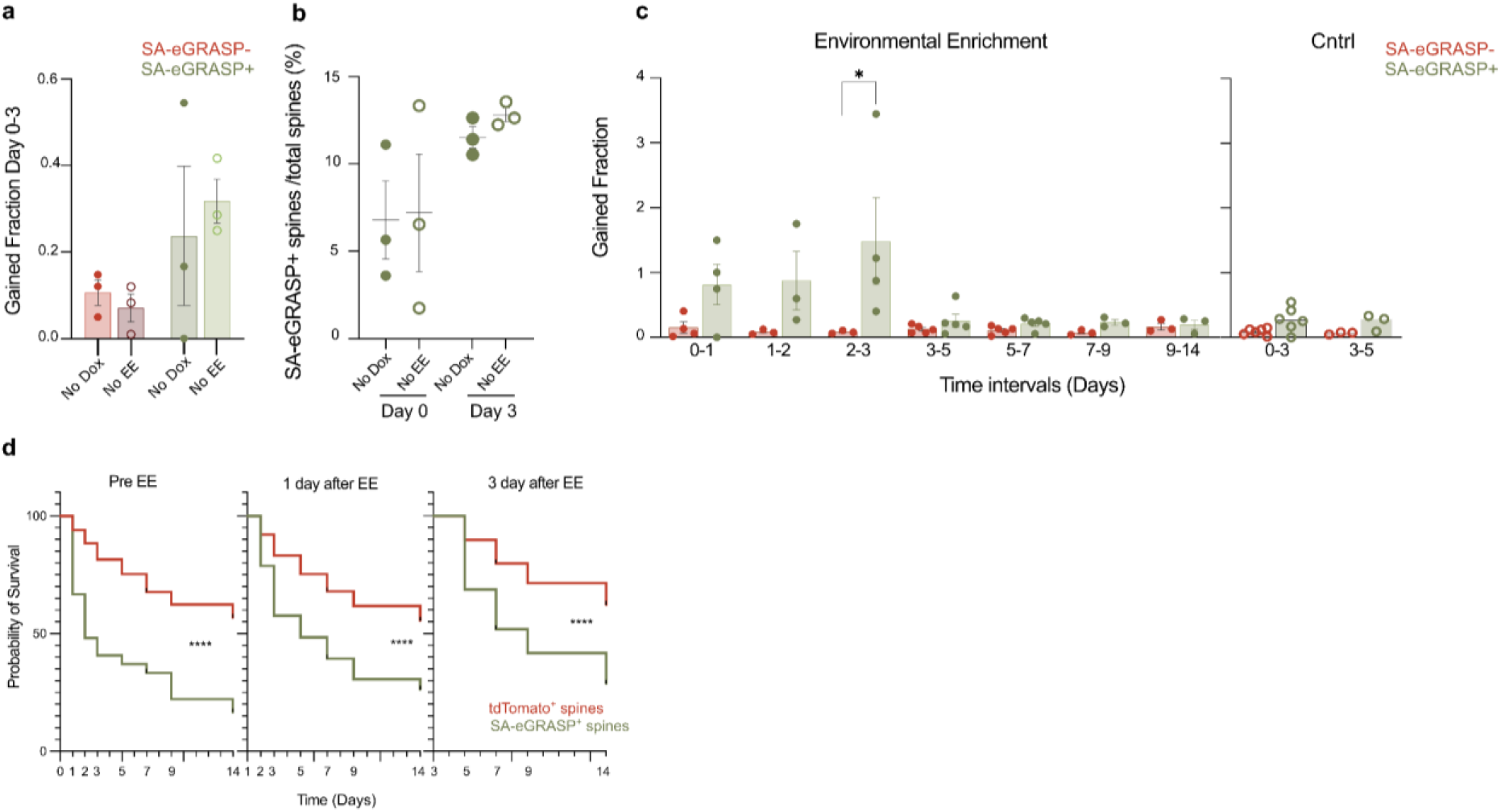
SA-eGRASP expression dynamics *in vivo*. **a.** Fraction of newly formed SA-eGRASP+ (green) and SA-eGRASP-(red) spines per mouse at day 0 and 3 in the two control groups (ANOVA-1, *F*_3,8_ = 1.729, *P* = 0.2380; each data point represents one mouse). **b**. Percentage of SA-eGRASP+ over all spines per mouse in Control groups (ANOVA-1, *F*_3,8_ = 2.186, *P* = 0.1675; each data point represents one mouse). **c**. Fraction of newly formed SA-eGRASP+ (green) and SA-eGRASP-(red) spines per mouse for both EE and Control groups (EE group, ANOVA-1 *F*_13,39_ = 3.105, *P* = 0.0031; Holm–Šídák post-hoc test, E vs. F, *P*_Int. 2-3 SA-eGRASP-vs SA-eGRASP+_ = 0.0236; control group, ANOVA-1, *F*_3,14_ = 3.191, *P* = 0.0566; each data point represents one mouse. **d**. Kaplan-Meier survival curves of SA-eGRASP+ puncta (green) and SA-eGRASP spines (red) across three reference time points: Day 0 (pre-EE), Day 1 (acute post-EE), and Day 3 (post-EE). For Day 0, Gehan–Breslow– Wilcoxon test: *χ²*_1_ = 33.89, *P* < 0.0001. SA-eGRASP+ puncta, *N* = 27 puncta (21 events, 6 censored); SA-eGRASP spine, *N* = 449 puncta (182 events, 267 censored). For Day 1, Gehan–Breslow–Wilcoxon test: *χ²*_1_ = 17.14, *P* < 0.0001, *N*_SA-eGRASP+ puncta_ = 33 puncta (23 events, 10 censored); *N*_SA-eGRASP+ spine_ = 453 puncta (193 events, 260 censored). For Day 3, Gehan–Breslow–Wilcoxon test: *χ²*_1_ = 43.59, *P* < 0.0001; *N*_SA-eGRASP+ puncta_ = 83 puncta (56 events, 27 censored); *N*_SA-eGRASP spine_ = 499 puncta (169 events, 330 censored). Data were obtained from 5 mice. Censored puncta remained present at the end of the observation period.

We imaged all mice one day before and at multiple time points following EE, for up to 14 days (Fig. 5b). During each imaging session, we anesthetized and head-fixed mice, then acquired three-dimensional, two-channel image stacks (green, SA-eGRASP; red, tdTomato) (Fig. 5c). We detected an average 93.24 tdTomato+ spines and 11.74 SA-eGRASP+ puncta localized on tdTomato+ spines in the EE group and 103.4 tdTomato+ spines and 9 SA-eGRASP+ puncta localized on tdTomato+ spines in the Control group, per mouse per day (Fig. 5d). The proportion of SA-GRASP^+^ spines at each time point varied between 2% and 25% of all spines with a significant increase in the EE over the control group at day 3 (Fig. 5d,e). In mice exposed to EE, the fraction of SA-eGRASP+ spines gained between consecutive imaging time points exceeded that of SA-eGRASP-spines, reaching statistical significance by day 3. In contrast, control animals exhibited no detectable increase in the fraction of SA-eGRASP+ spines gained (Extended Data Fig. 8c). Direct comparison of the same time intervals in the two groups confirmed an increase in the number of SA-eGRASP+ spines within 3 days after EE (Fig. 5f). SA-eGRASP+ puncta predominantly appeared on pre-existing spines throughout the experiment both in the EE and Control groups (Fig. 5g, h). Interestingly, however, exposure to EE also led to a trend towards an increase in the proportion of newly potentiated SA-eGRASP+ puncta appearing on newly formed spines (Fig. 5i).

Altogether, these data show an increase in SA-eGRASP+ dendritic spines in the dCA1, peaking 3 days after exposure to EE.

We then compared the long-term stability of SA-eGRASP+ and SA-eGRASP-spines, formed either before EE exposure (day 0), immediately after EE exposure (day 1), or following EE exposure (day 3). SA-eGRASP+ spines appearing on all days were significantly less persistent than SA-eGRASP-spines (Extended Data Fig. 8d). Among all observed SA-eGRASP^+^ spines, we found that in a minority of cases (∼20%) the SA-eGRASP signal was lost while the spine remained structurally intact; however, in most instances (∼80%), SA-eGRASP^+^ spines were pruned (Fig. 6a,b).

**Figure 6.**
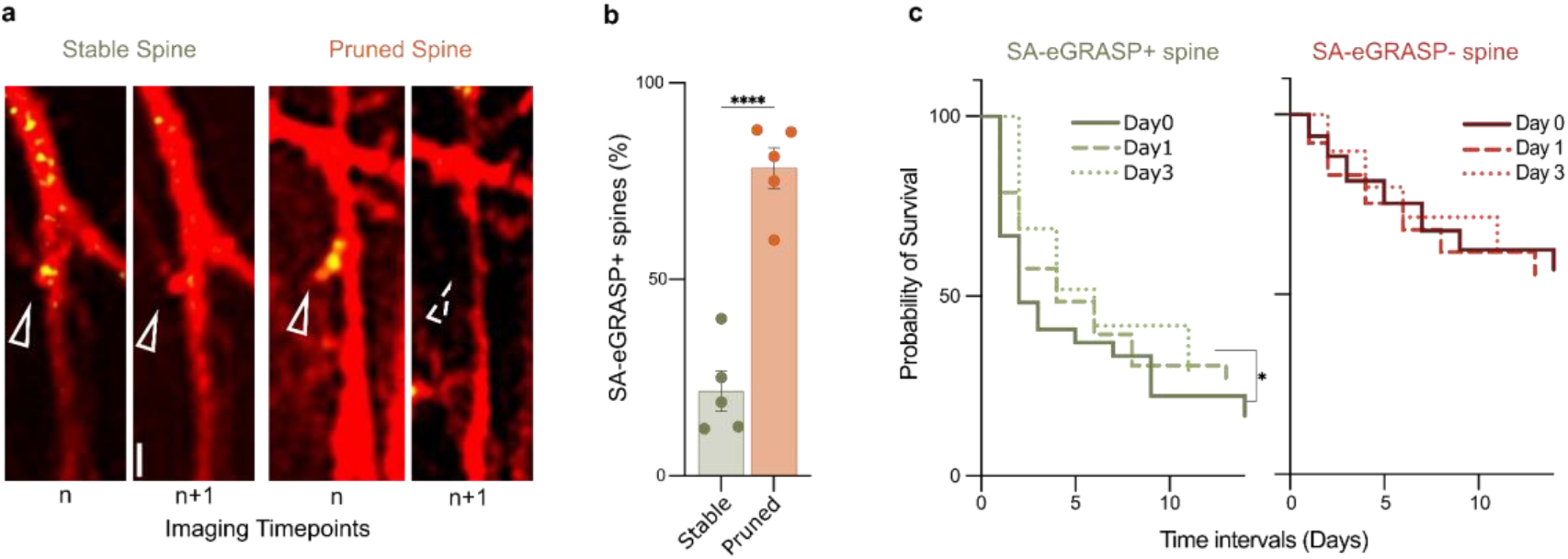
A novel experience leads to stabilization of SA-eGRASP+ spines over days. **a.** Representative 2P images (single z-plane) of SA-eGRASP puncta (green) that were eliminated on spines that remained stable (left) or were pruned (right) at the next imaging session. Scale bar, 2 μm. **b.** Percentage of SA-eGRASP+ puncta, EE group, on dendritic spines that were stable or were pruned in the next imaging session (*t*-test, *t*_8_ = 7.765, *P* < 0.0001). Each data point represents one mouse. **c**. Kaplan–Meier survival curves for SA-eGRASP+ puncta and SA-eGRASP-spines detected on Day 0 (pre-EE, full line), Day 1 (acute post-EE, dashed line), to Day 3 (post-EE, dotted line). Gehan–Breslow–Wilcoxon test, *χ²*_2 SA-eGRASP+_ = 6.710, *P* = 0.0349; *χ²*_2 SA-eGRASP-_ = 3.776, *P* = 0.1514. Uncorrected pairwise log-rank tests: SA-eGRASP+, *P*_Day0 vs. Day1_ = 0.5361, *P*_Day0 vs. Day3_ = 0.0387; N_Day0_ = 27 spines (6 censored), N_Day1_ = 33 spines (10 censored), N_Day3_ = 83 spines (27 censored); SA-eGRASP-, *P*_Day0 vs. Day1_ = 0.0781, *P*_Day0 vs. Day3_ = 0.6617, *P*_Day1 vs. Day3_ = 0.3246; N_Day1_ = 449 spines (182 events, 267 censored), N_Day1_ = 453 spines (193 events, 260 censored); N_Day3_ = 499 spines (169 events, 330 censored); N_mice_ = 5. Censored spines were those remaining present at the end of imaging.

These data indicate that dendritic spines expressing SA-eGRASP+ have a shorter lifetime than SA-eGRASP-spines.

We next asked whether exposure to EE leads to stabilization of SA-eGRASP+ spines. To this aim, we compared the survival of SA-eGRASP+ spines detected before EE (day 0) with that of those detected after EE exposure (day 1 and 3). SA-eGRASP+ spines detected on day 3, but not on day 1, were significantly more stable compared to those detected at day 0 (Fig. 6c). This effect was specific to SA-eGRASP+ spines and was not observed in SA-eGRASP-spines (Fig. 6c).

Together, these findings indicate that EE selectively stabilizes a subset of spines labeled by SA-eGRASP.

## DISCUSSION

Circuit-specific synaptic plasticity is thought to underlie information storage, yet its spatial and temporal organization across defined neuronal connections remain incompletely understood. Here, we developed SA-eGRASP, a genetically encoded reporter that combines activity-dependent postsynaptic labeling with trans-synaptic GFP reconstitution to identify potentiated synapses between defined neuronal populations *in vivo*. Using this approach, we mapped dCA3–dCA1 synaptic potentiation across distinct phases of associative memory formation and tracked its evolution longitudinally in live animals. Our findings reveal the circuit-, dendrite- and synapse-level organization of plasticity associated with memory formation and retrieval.

While existing tools allow capturing synaptic activity, receptor trafficking, or structural dynamics^25–30^, SA-based reporters selectively report potentiation at the synaptic level as a result of local synthesis and *via* the use of peptide sequences for localization at the synapse^16,17,20^. However, their spatial precision can be limited by the spread of the translational machinery in dendrites and synapses^31^. SA-eGRASP solves this problem by combining local postsynaptic activation with the requirement for a presynaptic connection. Thus, SA-eGRASP encodes a conditional *AND* logic gate, restricting labeling to individual synapses that are both structurally connected and functionally potentiated. Moreover, while the conventional GRASP system under the control of IEG promoters^3,18,19^ maps overall connectivity, without distinguishing the functional state of synapses, SA-eGRASP preferentially labels individual potentiated synapses *via* activity-dependent local synthesis. Hence, SA-eGRASP provides a closer approximation to functional synaptic modification and reduces the spatial ambiguity inherent to postsynaptic-only or activity-defined neuronal labeling strategies^5,13,15^.

SA-eGRASP labelling revealed several features of potentiated CA3-CA1 synapse remodeling associated with learning and recall of a hippocampus-dependent associative memory task, *i.e.* CFC. First, encoding increased both the density and volume of SA-eGRASP+ puncta, whereas recall increased puncta volume and altered their spatial distribution with respect to *naïve* mice (HC group). These findings are consistent with synaptic strengthening within hippocampal circuits^1,32,33^ and selective modification of memory engram-associated networks upon learning^5,13–15^. Second, the different topography of potentiated synapses after encoding and recall suggests that these processes engage distinct forms of plasticity. Specifically, the increase in volume of SA-eGRASP+ puncta upon recall — with no significant difference in their density or abundance — suggests selective engagement of synapses activated during encoding, thus reflecting consolidation of a subset of existing connections^34,35^, while others return to baseline or are pruned^36,37^. Importantly, we recently reported structural signatures of memory retrieval in a subset of previously activated synapses, consistent with the idea that recall preferentially recruits established connections rather than inducing widespread new potentiation^17^. Third, dendritic clustering of SA-eGRASP+ puncta is consistent with the local organization of synaptic plasticity. The recall-blank experiment indicates that SA-eGRASP labeling can resolve synaptic potentiation associated with retrieval independently of prior learning-induced labeling.

Whereas previous work described the anatomical distribution of synaptic inputs independent of their functional state^13^, we show that learning selectively engages groups of neighboring synapses, suggesting that synaptic clustering reflects not only circuit architecture but also local organization of plasticity^38^. This view is consistent with studies showing that dendritic branches act as functional units of integration and plasticity^39–42^, and with models proposing that synaptic clustering enhances storage capacity and supports nonlinear computation^43,44^. Notably, our data suggest that the identity of a neuron within an engram network is not binary — as IEG-based approaches to label engram neurons might imply — but that different branches within a neuron can contribute variably to information processing and storage by virtue of different patterns of synaptic potentiation. In these terms, SA-eGRASP provides a novel framework to elucidate the function of learning-related potentiation of specific synapses and how these are distributed along different dendritic segments within single neurons, irrespective of (and in addition to) their somatic IEG expression levels. In turn, this enables investigating synaptic engrams, in a way that has not been possible thus far.

The strong fluorescence and high signal-to-noise ratio of SA-eGRASP enabled us to track how individual dCA1-dCA3 activated synapses evolved. While CFC induced rapid and widespread SA-eGRASP expression, EE produced slower and sparser labeling, with delayed peak dynamics. These differences likely reflect both the strength and nature of the inducing experience, as well as methodological sensitivity and sample size.

While the SA-eGRASP signal mostly appeared on pre-existing spines, the proportion of SA-eGRASP+ new spines showed a trend towards an increase upon EE. This is consistent with previous studies showing that enhanced activity- and experience-dependent plasticity triggered by EE can lead to both strengthening of existing and formation of new synapses^13,45–47^, and suggests that activity-dependent local synthesis occurs preferentially at established synaptic contacts. Interestingly, SA-eGRASP+ spines were less stable than SA-eGRASP-ones, and the disappearance of SA-eGRASP strongly correlated with the disappearance of the spines on which these puncta were localized. This shows that recently potentiated spines are, on average, less stable than other spines. Together with the previous finding, this may imply that, while new synaptic engrams are preferentially allocated to existing synapses, turnover of synaptic engrams is associated with synapse and spine loss. This loss is presumably balanced by the constant appearance of new synapses^48–50^.

Notably, SA-eGRASP+ spines labelled upon EE showed enhanced persistence in comparison to SA-eGRASP+ spines labelled prior to EE. This shows that — despite the intrinsic instability of recently-potentiated spines — naturalistic neuronal activation leads to stabilization of a subset of specific synaptic contacts. Some caution should be taken while interpreting these results, as SA-eGRASP reports structural correlates of plasticity rather than direct functional potentiation^16,17,20^, and relies on Arc regulatory elements, which restrict detection to specific forms of plasticity^51^. Addressing these limitations will require integration with electrophysiological or optical activity measurements and systematic comparison with existing approaches, including IEG-driven^5,13–15^ and SA-based methods^16,17,20^.

Of note, SA-eGRASP can be further developed to simultaneously map synaptic potentiation at multiple circuits by employing, for instance, spectral variants of split GFP^14^ and also expanded to label inhibitory synapses, whose dynamics show a complex spatial pattern upon learning^52^.

Thus, SA-eGRASP provides a practical and scalable approach to identify and track potentiated synapses within defined circuits *in vivo*^3,5,15^. By enabling tracking of the same synapses over time, this method bridges an important gap between cellular engram studies and synapse-level investigation of memory^1,5,15^. More broadly, SA-eGRASP provides a foundational method that will shift our understanding of memory engrams from the whole-neuron to the single-synapse resolution.

## ACKNOWLEDGEMENTS

We gratefully acknowledge: Dr. Alessandro Viegi, Dr. Maria Antonietta Calvello, Mrs. Vania Liverani for technical, infrastructural, and organizational assistance; Francesca Biondi for animal colony care and maintenance; Prof. Cristina Alberini (NYU), Dr. Silvia Marinelli (EBRI), Dr. Alessandro Ulivi (University of Heidelberg), and Prof. Federico Cremisi (SNS) for scientific discussions; Dr. Andrea Faraone and Dr. Mariachiara Di Caprio for contributing to previous and parallel SynActive lab work; Mr. Tommaso Gosetti Di Sturmeck and Ms. Giorgia Ribbeni for their contribution to the ongoing expansion of the SynActive dataset, and production of the analysis user manual. We thank Prof. Alexander Dityatev’s lab (DZNE Magdeburg) and especially Ms. Katrin Boehm for the production and purification of AAVs employed in *in vivo* imaging experiments.

## AUTHOR CONTRIBUTIONS

FCL, investigation, formal analysis, writing-original draft, software, data curation, performed *in vivo* AAV injections, behavioral tests, *ex vivo* imaging experiments, contributed to MATLAB-based analysis tools, performed statistical analyses. AJ, investigation, formal analysis, writing-original draft, software, data curation, performed *in vitro* experiments, developed the MATLAB code, and performed statistical analyses. HM, conceptualization, investigation, formal analysis, data curation, designed and performed *in vivo* time-lapse experiments and analyzed data. AC, conceptualization, supervision, funding acquisition, writing-review & editing. AA, conceptualization, supervision, funding acquisition, writing-original draft, writing-review & editing, designed *in vivo* imaging experiments. MM, conceptualization, supervision, funding acquisition, investigation, writing-original draft, writing-review & editing, contributed to behavioral and *ex vivo* imaging experiments, provided conceptual and technical guidance. All authors discussed the results and reviewed and approved the final manuscript.

